# Reprogramming of the apoplast metabolome of *Lolium perenne* upon infection with the mutualistic symbiont *Epichloë festucae*

**DOI:** 10.1101/861450

**Authors:** Kimberly A Green, Daniel Berry, Kirstin Feussner, Carla J. Eaton, Arvina Ram, Carl H. Mesarich, Peter Solomon, Ivo Feussner, Barry Scott

## Abstract

*Epichloë festucae* is an endophytic fungus that forms a mutualistic symbiotic association with *Lolium perenne*. Here we analysed how the metabolome of the ryegrass apoplast changed upon infection of this host with sexual and asexual isolates of *E. festucae*. A metabolite fingerprinting approach was used to analyse the metabolite composition of apoplastic wash fluid from non-infected and infected *L. perenne*. Metabolites enriched or depleted in one or both of these treatments were identified using a set of interactive tools. A genetic approach in combination with tandem mass spectrometry was used to identify a novel product of a secondary metabolite gene cluster. Metabolites likely to be present in the apoplast were identified using the MarVis Pathway in combination with the BioCyc and KEGG databases, and an in-house *Epichloë* metabolite database. We were able to identify the known endophyte-specific metabolites, peramine and epichloëcyclins, as well as a large number of unknown markers. To determine whether these methods can be applied to the identification of novel *Epichloë*-derived metabolites, we deleted a gene encoding a NRPS (*lgsA*) that is highly expressed *in planta*. Comparative mass spectrometric analysis of apoplastic wash fluid from wild-type- versus mutant- infected plants identified a novel Leu/Ile glycoside metabolite present in the former.

## Introduction

The fungal endophyte *Epichloë festucae* forms symbiotic associations with temperate grasses of the *Festuca* and *Lolium* genera (Leuchtmann *et al*., 1994). Hyphae grow in the intercellular (apoplastic) spaces of aerial tissues, systemically colonising the host leaf sheath, leaf blade, and inflorescences (May *et al*., 2008; Scott *et al*., 2012). Hyphae attach to the host cell wall through an adhesive matrix, and cease growing in the later stages of host development, though they remain metabolically active (Tan *et al*., 2001; Christensen & Voissey, 2007). *Epichloë* endophytes growing *in planta* produce a wide variety of secondary metabolites (SMs) that protect the host against herbivores (Malinowski & Belesky, 2000; Clay & Schardl, 2002; Saikkonen *et al*., 2016). The most well characterized *Epichloë* SMs are the ergot alkaloids, lolines, indole-diterpenes, and pyrrolopyrazines such as peramine (Young *et al*., 2005; Schardl *et al*., 2013; Berry *et al*., 2015). Ergot alkaloids and indole diterpenes are anti-mammalian mycotoxins (Florea *et al*., 2016; Philippe, 2016), while lolines and peramine protect against insect herbivores (Rowan & Gaynor, 1986; Wilkinson *et al*., 2000; Tanaka *et al*., 2005; Schardl *et al*., 2007; Pan *et al*., 2014).

In addition to these well-known SMs, *Epichloë* endophytes are known to produce ribosomally synthesized and post-translationally modified peptides (RiPPs) derived from the secreted polypeptide GigA (Johnson *et al*., 2015). Imperfect 27-amino acid repeats within the GigA sequence are cleaved to release multiple different cyclic RiPPs, which are collectively known as the “epichloëcyclins”. The spectrum of epichloëcyclins produced varies between different endophyte strains due to differences in encoded GigA sequences; for example, epichloëcyclins A–N have been characterized, but only A–E are present in *E. festucae* strain Fl1 (Johnson *et al*., 2015). *Epichloë* species are also known to protect the host, to varying degrees, from a range of fungal pathogens (Gwinn & Gavin, 1992; Bonos *et al*., 2005; Clarke *et al*., 2006; Steinebrunner *et al*., 2008; Niones & Takemoto, 2014; Niones & Takemoto, 2015; Tian *et al*., 2017). In addition to protection from these biotic stresses, *Epichloë* endophytes also protect the host from abiotic stresses such as drought (Arachevaleta *et al*., 1989; Hahn *et al*., 2007) although the physiological basis of this phenomenon is not known. Collectively these, and other studies, have shown that *Epichloë* endophytes have an important biological role in both natural and agricultural ecosystems (Johnson *et al*., 2013).

There are many changes in the host transcriptome upon endophyte infection including differential expression of genes involved in host hormone production, pathogenesis-related gene expression, adaptation to biotic stresses, and host cell wall metabolism (Ambrose & Belanger, 2012; Dupont *et al*., 2015; Schmid *et al*., 2016; Dinkins *et al*., 2017). Given these major changes in the transcriptome it is likely that endophyte infection results in metabolome reprogramming. While transcriptome studies have given us significant insights into how host and endophyte reprogramming might occur (Eaton *et al*., 2010; Eaton *et al*., 2015; Chujo *et al*., 2019), these experiments provide very little insight into the exact metabolite changes that occur within the apoplastic environment where the two organisms interact. While some of the SMs synthesized by *Epichloë* are known to be mobilised throughout the host plant, presumably facilitating systemic host protection (Koulman *et al*., 2007), little is known about their exact composition in the apoplast. The aim of this study was to compare the composition of metabolites in the apoplast wash fluid of *Epichloë*-infected vs. mock-infected plants by a non-targeted liquid chromatography-coupled high-resolution electrospray ionization mass spectroscopy (LC-HR-MS)-based metabolome analysis. In particular, we set out to identify unknown compounds that could be novel infection-induced *E. festucae* or host synthesised metabolites that contribute to host protection.

## Materials and Methods

### Endophyte inoculations and growth conditions

Endophyte-free and common-toxic (CT)-infected seeds (*Lolium perenne* cv. Samson) were germinated on 3% water agar. Endophyte-free seedlings were inoculated with *E. festucae* strains as previously described (Latch & Christensen, 1985). Plants were grown in root trainers in an environmentally controlled growth room at 22°C with a photoperiod of 16 h of light (∼100 μE/m^2^ per sec) and at 10 weeks post-inoculation, tested for the presence of the endophyte by immunoblotting (Tanaka *et al*., 2005).

Two-week old seedlings of *Triticum aestivum* (*cv* Summit) were infected with *Zymoseptoria tritici* (WAI321) by syringe infiltration into the second leaf with spores at a concentration of 5×10^6^ mL^-1^.

*Escherichia coli* cultures were grown in lysogeny broth (LB) or on LB agar supplemented with 100 μg/mL ampicillin (Miller, 1972). *E. festucae* cultures were grown on 2.4% (w/v) potato-dextrose (PD) agar or in PD broth (Moon *et al*., 1999; Moon *et al*., 2000).

### Generation of gene deletion and complementation strains

Biological materials and primers can be found in Supporting Information Tables **S1** and **S2**. Plasmid DNA was extracted using the High Pure Plasmid Isolation Kit (Roche, Basel, Switzerland). Fungal DNA was extracted as previously described (Byrd *et al*., 1990). Cloning and screening PCR reactions were performed using Q5 High-Fidelity DNA and One-*Taq* DNA polymerases (New England Biolabs (NEB), Ipswich, USA) respectively. Products of PCR reactions were purified using the Wizard^®^ SV Gel and PCR Clean-Up system (Promega, Madison, USA). Sequencing reactions were performed using the Big-DyeTM Terminator Version 3.1 Ready Reaction Cycle Sequencing Kit (Applied BioSystems, Carlsbad, California, USA), and separated using an ABI3730 genetic analyser (Applied Bio Systems, Carlsbad, California, USA). Sequence data was assembled and analysed using MacVector sequence assembly software, version 12.0.5 (MacVector Inc., Apex, NC, USA).

**Table 1.** Putative functions of *LGS* cluster-proximal genes in *Epichloë*, *Zymoseptoria*, and *Ramularia* spp.

| Gene | Gene model(s)<sup>1</sup> | Putative function |
| --- | --- | --- |
| *lgsA* | EfM3.056230,<br>ZT1E4_G1886,<br>TI39_contig403g00012,<br>RCC_03904 | NRPS-like, A-T-C domain structure |
| *lgsB* | EfM3.062330,<br>ZT1E4_G1887,<br>TI39_contig42g00014 | NADP-binding oxidoreductase |
| *lgsC* | EfM3.062310 | glycosyl transferase type 2 |
| *lgsD* | EfM3.056220 | ABC transporter type I |
| 1 | EfM3.056240 | haloacid dehydrogenase-like hydrolase |
| 2 | EfM3.062340 | mitochondrial substrate carrier |
| 3 | EfM3.062320 | FAD-binding oxidoreductase |
| 4 | ZT1E4_G1891,<br>TI39_contig42g00011 | transcription factor |
| 5 | ZT1E4_G1890,<br>TI39_contig42g00012 | MFS sugar transporter |
| 6 | ZT1E4_G1889,<br>TI39_contig42g00010 | anhydro-N-acetylmuramic acid kinase |
| 7 | ZT1E4_G1888,<br>TI39_contig42g00013 | DNA repair metallo-beta-lactamase |
| 8 | ZT1E4_G1885,<br>TI39_contig403g00011 | Integral membrane protein, unknown function |
| 9 | ZT1E4_G1884,<br>TI39_contig403g00010 | nitrogen permease regulator 3 |
| 10 | ZT1E4_G1883,<br>TI39_contig403g00009 | mitochondrial substrate carrier |
| 11 | ZT1E4_G1882,<br>TI39_contig403g00008 | integral membrane protein, collagen binding/endopeptidase activity |
| 12 | RCC_03900 | Ubiquitin-conjugating enzyme E2 |
| 13 | RCC_03901 | Integral membrane protein, SUR7/Rim9-like |
| 14 | RCC_03902 | unknown |
| 15 | RCC_03903 | histidine phosphatase superfamily |
| 16 | RCC_03905 | glycoside hydrolase family 47 |
| 17 | RCC_03906 | sodium/calcium exchanger |
| 18 | RCC_03907 | amino acid/polyamine transport |
| 19 | RCC_03908 | unknown |

<sup>1</sup> EfM3 gene models for *E. festucae* are from the UKY genome database (http://www.endophyte.uky.edu). Gene models for *Z. tritici* (ZT1E4), *Z. brevis* (TI39), and *R. collo-cygni* (RCC) are from the Ensembl Fungi database (http://fungi.ensembl.org/index.html).

The *lgsA* deletion and complementation constructs were assembled using Gibson Assembly (Gibson *et al*., 2009). The *lgsA* (gene model EfM3.028480) (Schardl *et al*., 2013) replacement construct pKG39 was prepared by recombining a 2.6 kb pAN7-1 vector backbone (amplified using primers pRS426F/pRS426R from a pAN7-1 plasmid DNA template), a 1.4 kb hygromycin resistance cassette (primers hphF/hphR; pSF15.15 plasmid DNA template), and 1 kb and 2.3 kb sequence fragments that flank the *lgsA* gene in the Fl1 genome (primers KG167/168 and KG169/170, respectively; *E. festucae* Fl1 genomic DNA template). The Δ*lgsA* complementation construct pKG40 was prepared using a 5.5 kb pRS426 backbone (primers pRS426F/pRS426R; pRS426 plasmid DNA template) and a 4.2 kb genomic DNA fragment containing *lgsA* (primers KG167KG170; *E. festucae* Fl1 genomic DNA template). Plasmids were transformed into chemically competent *E. coli* DH5α cells and transformants selected using ampicillin (100 μg/mL), and screened by restriction enzyme digestion. Candidate clones were verified by insert sequencing.

The *lgsC* (originally named *cpsA;* gene model EfM3.028480) deletion construct pCE57 was assembled using yeast recombinational cloning (Colot *et al*., 2006). For this, 5’ and 3’ sequences flanking the *lgsC* gene of 1432 bp and 967 bp respectively, were PCR-amplified using Phusion^®^ polymerase from genomic DNA template using primer sets pRS426-cpsA-F/cpsA-hph-R and hph-cpsA-F/cpsA-pRS426-R, which contained sequences that overlap with the yeast vector pRS426 and hph cassette. The P*trpC-hph* cassette was amplified with primers hph-F/hphR. Yeast cells were transformed with the *Eco*RI/*Xho*I-linearised pRS426 backbone together with the *lgsC* 5’ and 3’ flanks and the P*trpC-hph* cassette, as previously described (Gietz & Woods, 2002). Transformants were selected on media lacking uracil, and plasmid DNA isolated and transformed into *E. coli* to select for plasmids containing the *in silico-* predicted sequence of pCE57.

*E. festucae* protoplasts (Young *et al*., 2005) were prepared and transformed with 2-3 μg of target DNA, as previously described (Itoh *et al*., 1994). *E. festucae lgsA* deletion strains were obtained using the Split Marker system (Rahnama *et al*., 2016). Two PCR fragments (primers KG171/SplitR and SplitF/KG172, pKG39 plasmid DNA template) were transformed into Fl1 protoplasts and transformants selected using hygromycin (150 μg/mL). *E. festucae lgsC* deletion strains were obtained by transforming a PCR-amplified linear product from pCE57 using primers AR84/AR85. Transformants were sub-cultured three times onto selection media and putative *lgsA* ‘knockouts’ were PCR-screened using primer sets KG173/174 (1.5 kb knock-out and 1.1 kb wild-type (WT) products), and KG175/10 and KG11/176 (1.2 kb and 2.3 kb products for 5’ and 3’ flanks where correct integration has occurred). *lgsC* putative‘ knockouts’ were PCR-screened using primer sets cps7/cps8 (4 kb knock-out and 3.7 kb WT products), and cps7/TC44 and cps8/TC45 (1.56 kb and 2.04 kb products for 5’ and 3’ flanks where correct integration has occurred).

Candidate ‘knockouts’ were confirmed by Southern analysis (Southern, 1975). Genomic DNA restriction enzyme digests (New England Biolabs (NEB), Ipswich, USA), separated by agarose gel electrophoresis, were transferred to positively charged nylon membranes (Roche, Basel, Switzerland) and fixed by UV light cross-linking in a Cex-800 UV light cross-linker (Ultra-Lum, Claremont, California, USA) at 254 nm for 2 min. Digoxigenin-dUTP (DIG) labelling, and hybridisation of the pKG39 DNA probe, and nitroblue tetrazolium chloride and 5-bromo-4-chloro-3-indolyl-phosphate (NBT/BCIP) visualisation, were performed using the DIG High Prime DNA Labelling and Detection Starter Kit I (Roche, Basel, Switzerland). *Hin*dIII produces 8 kb, 2.6 kb and 1.8 kb fragments in WT, and 10.9 kb and 1.8 kb fragments in ‘knockout’ strains. Complementation strains were generated by co-transforming pKG40 and pII99 plasmid DNA into Δ*lgsA* protoplasts using geneticin (200 μg/mL) selection. Complementation strains were PCR-screened using the primers KG171/172 (4.2 kb complementation product).

### Microscopy analyses

Culture morphology was analysed using an Olympus IX71 inverted fluorescence microscope, with the filters set for capturing DIC. Infected pseudostem tissues for Confocal Laser-Scanning Microscopy (CLSM) were stained as previously described (Becker *et al*., 2016; Becker *et al*., 2018). CLSM images were captured using a Leica SP5 DM6000B confocal microscope (488 nm argon and 561 nm DPSS laser, 40× oil immersion objective, NA=1.3) (Leica Microsystems, Wetzlar, Germany).

### Apoplast extractions for metabolite analysis

To extract apoplastic was fluid, *L. perenne* leaf and pseudostem tissues were rinsed and submerged in water, and vacuum-infiltrated at approx. 60 mbar for 3× 10 min cycles. Tissues were lightly blotted dry to remove surface water and placed in 50 mL polypropylene Falcon tubes retrofitted with a plastic sieve wedged above the conical section of the tube. Tubes were centrifuged at 2000 × ***g*** for 10 min to collect apoplastic wash fluid in the bottom of the tube, which was removed to a separate microtube and centrifuged at 17,000 × ***g*** for 10 min to remove cellular debris, then analysed or stored at -20°C.

Apoplastic wash fluid from infected and uninfected seedlings of *Triticum aestivum* (*cv* Summit) were harvested two weeks post-infection with *Z. tritici* (WAI321). The leaves were removed from the plant and submerged in sterile distilled water under vacuum for 5 min. The infiltrated leaves were then dried and placed into the barrel of a 30 mL syringe which in turn was inserted inside a sterile 50 mL centrifuge tube. The leaves were centrifuged at 5000 × ***g*** for 5 min at 4°C after which the extracted apoplast wash fluid was quickly transferred to a 1.5 mL Eppendorf tube and frozen in liquid nitrogen prior to being freeze-dried overnight and stored at -20°C.

### Metabolite fingerprinting

#### Two-phase extraction of apoplastic fluids

The two-phase extraction method with methyl-*tert*-butyl ether (MTBE), methanol and water (Matyash *et al*., 2008) was adapted from a previous method (Feussner & Feussner, 2019) and modified for apoplastic wash fluid. Frozen apoplastic wash fluid (-80°C) was thawed on ice, vortexed and 200 µl added to 150 µl of methanol in a glass vial followed by addition of 500 µl MTBE. The sample was vortexed and shaken for 1 h in the dark then 120 µl of water added. Samples were incubated for 10 min for phase separation and centrifuged for 15 min at 800 × ***g***. The upper (non-polar) and lower (polar) phases were separately removed, transferred to new glass vials, and dried under a stream of nitrogen. The metabolites of the polar extraction phase were dissolved in 60 µl of methanol/acetonitrile/water (1:1:12, *v/v/v*). The metabolites of the non-polar extraction phase were dissolved in 60 µl of 80% methanol. All samples were shaken for 15 min and centrifuged for 10 min. From each sample, 50 µl was transferred into a glass micro vial, covered with argon and used for metabolite fingerprinting analysis.

#### Data acquisition

For metabolite fingerprinting analysis an Ultra Performance Liquid Chromatography (UPLC) system coupled to the photo diode array (PDA) detector eλ (Waters Corporation, Milford, MA, USA) and the high resolution orthogonal Time-of-Flight Mass Spectrometer (HR-MS) LCT Premier (Waters Corporation, Milford, MA, USA) was used as previously described (Feussner & Feussner, 2019). Samples of the polar and the non-polar extraction phases were analysed in the positive as well as in the negative electrospray ionisation (ESI) mode, resulting in four data sets.

#### Data processing and data mining

Peak selection and alignment for each data set was done with the software MarkerLynx XS for MassLynx V.4.1 (Waters Corporation, Milford, MA, USA). This data deconvolution procedure resulted in four data matrixes of several thousand metabolite features (3258 and 1797 features for the polar extraction phase for positive and negative ESI respectively; 1750 and 740 features for the non-polar extraction phase for positive and negative ESI respectively (Supporting Information Tables **S3-S6**). The software suite MarVis (Kaever *et al*., 2015) (http://marvis.gobics.de/) was used for further data processing, such as ranking, filtering, adduct correction, merging of data sets as well as for data mining such as clustering and automated data base search. Subsequently, ANOVA and Benjamini-Hochberg for multiple testing were applied with the tool MarVis Filter to rank the metabolite features of each of the four data matrixes and filter them for a false discovery rates (FDR) of <0.003. After adduct correction the data matrixes were combined. This resulted in a data set of 203 metabolite features (Supporting Information Tables **S7** & **S8**). Next, the tool MarVis-Cluster was used for clustering and visualising the intensity profiles of the selected features by means of one-dimensional self-organizing maps (1D-SOMs). Finally, MarVis-Pathway was applied to facilitate the annotation of metabolite features based on accurate mass information. The databases KEGG (http://www.kegg.jp) (Tanabe & Kanehisa, 2012), BioCyc (http://biocyc.org) (Caspi *et al*., 2012) and an in-house database, specific for SMs of *Epichloë* (Supporting Information Table **S9**), were used in combination with a framework for metabolite set enrichment analysis to annotate the metabolite features.

#### Verification of the chemical structure of marker compounds

To confirm or to elucidate the chemical identity of marker metabolites, high resolution tandem mass spectrometry analyses (LC-HR-MSMS) were performed by UHPLC LC 1290 Infinity (Agilent Technologies, Santa Clara, CA, USA) coupled to the 6540 UHD Accurate-Mass Q-TOF LC-MS instrument with Agilent Dual Jet Stream Technology as ESI source (Agilent Technologies, Santa Clara, CA, USA).

### Whole-pseudostem extractions for polar metabolite analysis

Pseudostem tissues were snap-frozen in liquid nitrogen, freeze-dried, and 50 mg (dry weight) of each sample was placed into 2 mL impact-resistant screw-cap microtubes containing 3× 3.2 mm diameter stainless steel beads and ground to a powder using a FastPrep® FP120 Cell Disrupter System (Thermo Savant). Ground samples were mixed with 1 mL 50% (v/v) methanol and incubated at room temperature in the dark with end-over-end rotation at 40 rpm for 1 h. Cellular debris was removed by centrifugation at 17,000 × ***g*** for 10 min, and the supernatant transferred to an amber glass HPLC vial through a 13 mm diameter, 0.45-µm-pore polytetrafluoroethylene syringe filter (Jet BioFil, Guangzhou, China). Extracts were then analysed or stored at -20°C.

Whole pseudostem and apoplastic was fluid extracts generated for identification of the LgsA product were analysed using a Q Exactive™ Focus hybrid quadrupole-orbitrap high-resolution mass spectrometer (Thermo Fisher Scientific, Waltham, MA, USA). Each sample (5 µl injection) was separated on a 2.1× 150 mm ZORBAX Eclipse Plus C18 column with 1.8 µm particle size (Agilent Technologies, Santa Clara, CA, USA) with a flow rate of 0.2 mL/min and a linear gradient profile using water with 0.1% formic acid as eluent A and acetonitrile with 0.1% formic acid as eluent B, with time 0 min (*T*_0_) at 5% B, *T*_1_ at 5% B, *T*_21_ at 95% B, *T*_26_ at 95% B, *T*_27_ at 5% B, followed by equilibration to initial conditions over the following 6 min.

Samples (five replicates) were analysed by high-pressure liquid chromatography-coupled HR positive ESI MS (LC-MS) using a capture window of 133-2000 *m*/*z*, and the resulting mass spectra were compared using Compound Discoverer 2.1 (Thermo Scientific, Waltham, MA, USA) to identify features exhibiting highly differential profiles between sample conditions. Mass spectra from samples of mock-inoculated (uninfected) plants were used as controls for baseline subtraction.

### Bioinformatics

The *E. festucae* WT (Fl1/E894) genome is available at http://csbio-l.csr.uky.edu/ef894-2011/ (Schardl *et al*., 2013). The WT axenic culture and *in planta* transcriptome data used here is available from the Sequence Read Archive (SRA) under Bioproject PPRJNA447872 (Hassing *et al*., 2019).

## Results

### Endophyte infection alters the metabolite composition of the host apoplast

In nature, the SM composition of *E. festucae*-infected host tissues varies considerably depending on the *E. festucae* strain used (Schardl *et al*., 2007; Young *et al*., 2009; Schardl *et al*., 2013). We therefore included two *E. festucae* strains, Fl1 and CT, in our analysis. The former is a commonly used laboratory strain that forms a stable association with *L. perenne* (Leuchtmann *et al*., 1994; Scott *et al*., 2012) while the latter is a naturally occurring asexual endophyte of *L. perenne* (Christensen *et al*., 1993). While a range of SMs are known to be synthesised in each of these associations, our knowledge of the biosynthetic capability of Fl1 is much more extensive because of the availability of a complete genome sequence and the relative ease with which mutants can be generated (Scott *et al*., 2012; Schardl *et al*., 2013; Winter *et al*., 2018). To determine how Fl1 and CT strains affect the host apoplastic metabolome, we extracted apoplastic wash fluid from leaves of mock-inoculated (Mock), Fl1-infected, and CT-infected *L. perenne* associations by two-phase extraction and analysed both extraction phases by a metabolite fingerprinting approach using LC-HR-MS in both positive and negative ESI-modes (Feussner & Feussner, 2019). This analysis resulted in several thousand metabolite features (Supporting Information Tables **S3-S6**). Filtering features of the four data sets by FDR <0.003, selected 203 metabolite features. Since LC-ESI-MS analyses tend to form adducts, these 203 features may represent in total about 60-80 metabolites, which show strong differences in their intensity profiles between treatments. Principal component analysis of this data set showed a clear separation on principal component 1 (PC1) for the apoplast metabolome of mock-treated and FI1-infected ryegrass, while PC2 separates all three treatments (Fig. **S1**). Next, the intensity profiles of the selected features were clustered and visualised by means of 1D-SOMs and organised into four clusters (Fig. **1a** and Supporting Information Table **S7**). Cluster 1 contains 36 metabolite features that are specifically enriched in CT-infected samples, cluster 3 contains 106 features that are specifically enriched in Fl1-infected samples, and cluster 4 contains 42 features that are enriched in both CT and Fl1-infected samples. Cluster 2 contains 19 metabolite features that are depleted in the apoplastic wash fluid of endophyte-infected plants in comparison to the mock-treated plants.

**Fig. 1.**
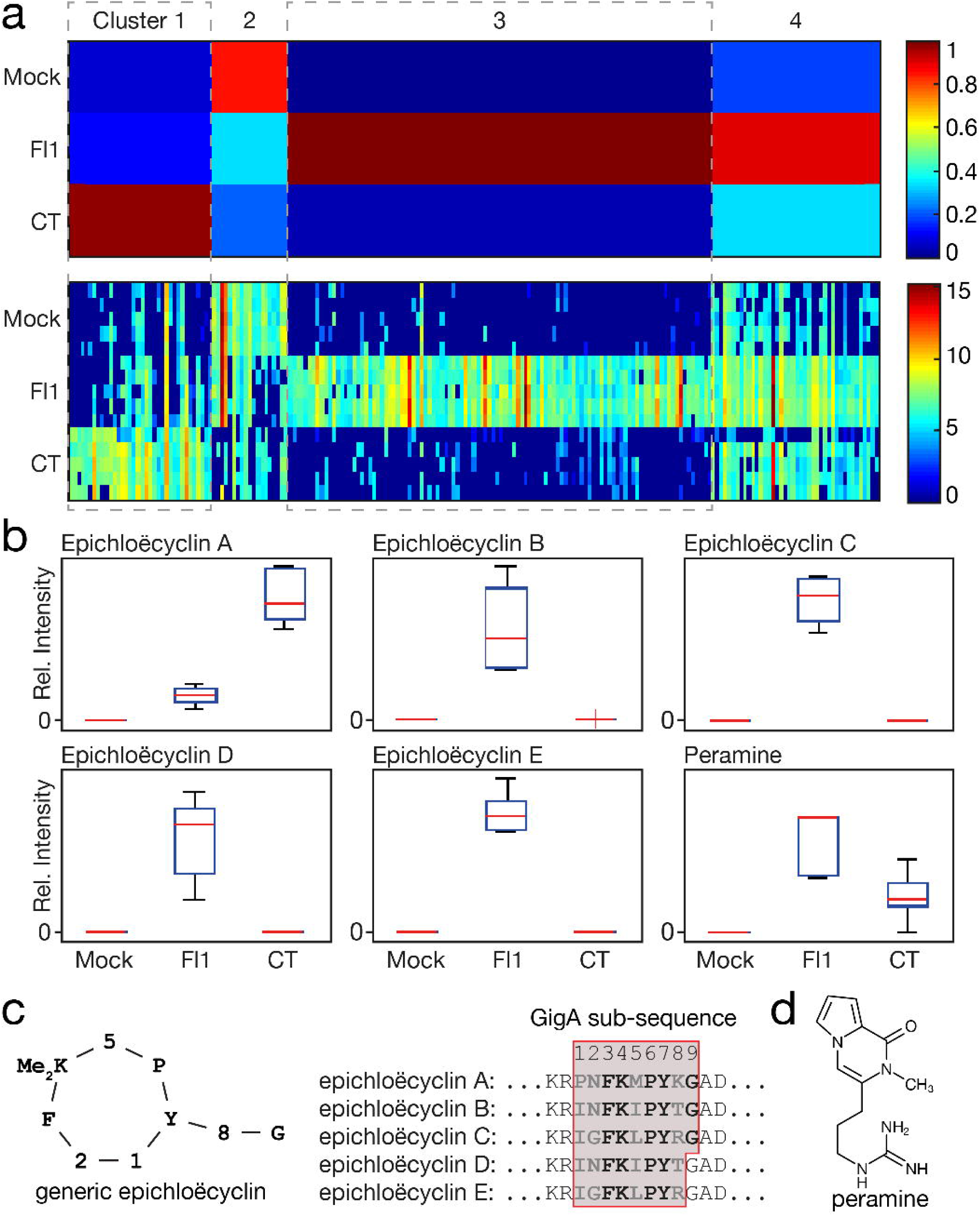
Metabolic fingerprinting of apoplastic wash fluid extracted from *E. festucae*-infected and non-infected *L. perenne*. Apoplastic wash fluid obtained from Mock-inoculated (M), Fl1-infected (F) and CT-infected (C) *L. perenne* associations at 18 weeks post-inoculation were analyzed by UPLC-ESI/QTOF-MS. (a) MarVis 1D-SOM intensity-based clustering of the 203 features (FDR < 0.003) detected in apoplastic wash fluid using ESI (blue low, red high). Five samples of M, F and C were analyzed. Polar and non-polar, positive and negative ESI data have been combined. (b) Metabolites confirmed to be in the apoplastic wash fluid samples by UHPLC-QTOF-HR-MSMS and their respective intensities. (c) Generic representation of epichloëcyclin structure and GigA repeat sequences from which epichloëcyclin A-E are derived. (d) Structure of peramine.

To assign identities to these features, the accurate mass information was used for a metabolite set enrichment analysis supported by the software tool MarVis Pathway (Kaever *et al*., 2015). The databases BioCyc and KEGG, as well as an in-house database, containing 117 SMs previously described for *Epichloë* (Supporting Information Table **S9**), were used to assign putative metabolite identities for these apoplastic features. The *E. festucae* metabolite peramine was identified in both CT and Fl1-infected samples, with a stronger accumulation in the Fl1-infected samples (Figs. **1b** & **1d**). The identity of peramine was confirmed unequivocally by LC-HR-MSMS analysis (Supporting Information Table **S8**; Fig. **S2**). A set of epichloëcyclins were detected as double charged ions ([M+2H]^2+^) using positive-mode ESI on *Epichloë*-infected samples (Figs. **1b** & **1c**; Supporting Information Table **S8**). Epichloëcyclin A was detected weakly in Fl1-infected samples and strongly in CT-infected samples, whereas epichloëcyclins B-E accumulated exclusively in Fl1-infected samples (Figs. **1b**, **S3** & **S4**). Sequencing *gigA* from CT, which is part of a four-gene cluster (Eaton *et al*., 2010) (Fig. **S5a**), revealed that this gene encodes a polypeptide that contains two identical epichloëcyclin A repeats (Fig. **S5b**). In addition to peramine and the epichloëcyclins, several other features representing so far unknown chemical structures were also present exclusively in the apoplastic wash fluid of endophyte-infected associations (Supporting Information Tables **S7** & **S8**) including two putative peptides, which are represented by the CT-specific cluster 1 (Fig. **S6**). A putatively annotated Kojibiose-related metabolite (322.0643 Da), a putatively annotated *N*-(hydroxypentyl) acetamide (145.1101 Da) and a compound of 471.1952 Da were found in the FI1-specific cluster 3 (Supporting Information Table **S8**). Such markers are of considerable interest as they represent potential novel bioactive fungal secondary metabolites.

### Identification of a novel amino acid glycoside produced by the NRPS-like protein LgsA

To determine whether rational biosynthetic predictions could be used to connect the novel metabolites identified by the non-targeted apoplast metabolome analysis to uncharacterized gene clusters, we chose to delete a gene (model EfM3.056230) encoding a non-ribosomal peptide synthetase (NRPS)-like protein that is part of a putative SM gene cluster (Fig. **2**) (Eaton *et al*., 2015). The genes from this cluster are expressed at very low levels in axenic cultures of *E. festucae* Fl1, but are dramatically upregulated when *E. festucae* is growing *in planta* (Fig. **2**; Supporting Information Tables **S10**) (Winter *et al*., 2018; Hassing *et al*., 2019). These genes were previously shown to be significantly down-regulated in three *L. perenne* associations containing symbiotically defective *E. festucae* mutants (Eaton *et al*., 2015). We have provisionally named this cluster *LGS* (<u>L</u>eucine/isoleucine <u>G</u>lycoside <u>S</u>ynthesis) based on the identification of the product described below. The NRPS-encoding gene (*lgsA*) from this cluster encodes the 1049 amino acid-long protein LgsA which contains an N-terminal thiolation (T) domain, a central condensation (C) domain, and a C-terminal adenylylation (A) domain (Fig. **S7**). The presence of an A-domain in LgsA suggests incorporation of an amino acid substrate into any product, but the domain structure of LgsA (T-C-A) is atypical compared to previously characterised NRPS and NRPS-like proteins. The other genes in the *LGS* cluster are predicted to encode an ABC transporter (*lgsD*), a haloacid dehydrogenase-like hydrolase (gene 1), a mitochondrial substrate carrier protein (gene 2), a NADP-binding oxidoreductase (*lgsB*), a FAD-dependent oxidoreductase (gene 3), and a glycosyl transferase (*lgsC*) (Table **1**). These putative functions suggested a product containing both amino acid and sugar residues that may be exported into the apoplastic space. To investigate the function of LgsA in *E. festucae*, a replacement construct (pKG39), was prepared and two overlapping PCR-amplified linear fragments of this plasmid were introduced into the genome of *E. festucae* strain Fl1 by homologous recombination (Fig. **S8**). PCR screening of ∼20 hygromycin-resistant transformants identified two candidates – Δ*lgsA*#2 and Δ*lgsA*#11 – that had PCR product patterns consistent with targeted replacement events. Southern analysis of genomic DNA digests from these transformants probed with a PCR fragment derived from pKG39 confirmed that both candidates contained a single-copy integration of the replacement construct at the target *lgsA* gene locus, with no additional ectopic integrations (Fig. **S8**).

**Fig. 2.**
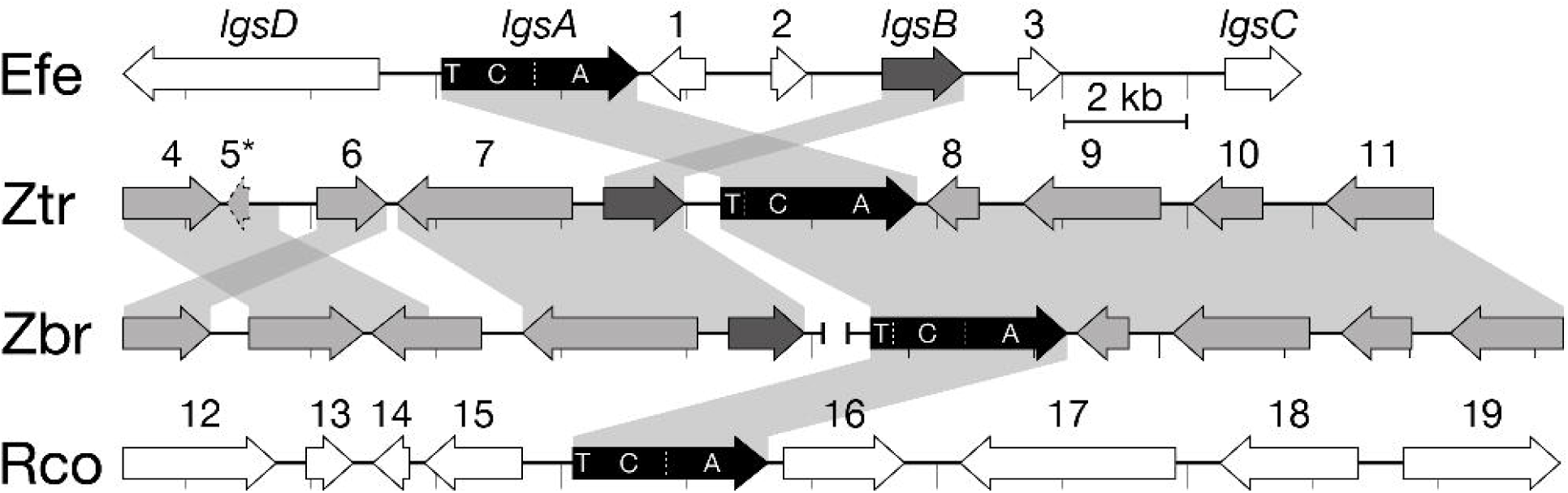
Microsyntenic comparison of *LGS* loci between *Epichloë*, *Zymoseptoria* and *Ramularia* spp. The *E. festucae LGS* cluster genes with known or suspected functions are named *lgsA*-*lgsC*, with all other genes assigned numbers. Predicted functions of all labelled genes are listed in Table 1. Syntenic blocks are indicated by grey shading, and darker gene colouring indicates a higher level of homology between species.

Macroscopic and microscopic analyses revealed that the culture morphologies of both Δ*lgsA* strains were indistinguishable from WT (Fig. **S9**). To determine the host interaction phenotype, WT and Δ*lgsA* strains were inoculated into *L. perenne* seedlings and the phenotype of infected plants was examined eight weeks post-planting (Fig. **S10**). The whole plant and cellular phenotypes of plants infected with the Δ*lgsA* mutants were indistinguishable from WT-infected plants (Fig. **S10**). These results were not unexpected given the likely function of the product of this gene is the synthesis of a bioprotective metabolite, which are typically phenotypically neutral when plants are grown under laboratory conditions in the absence of biotic stress (Tanaka *et al*., 2005; Young *et al*., 2005; Saikia *et al*., 2012).

To examine the chemotype of these associations whole-pseudostem extracts and apoplastic wash fluid samples were extracted from *L. perenne* plants infected with WT, Δ*lgsA* mutants (both Δ*lgsA*#2 & Δ*lgsA*#11), or from mock-inoculated plants. Comparison of the LC-MS data of pseudostem extract samples from Fl1-infected and Δ*lgsA*-infected plants did not identify any significantly different signal abundances. In contrast, comparison of apoplastic wash fluid samples between these conditions identified 16 significantly different features (P<0.05), with five of these features exhibiting the largest reduction in intensity in the Δ*lgsA* samples, as would be expected for a LGS metabolite. Four of these features were still present in the Δ*lgsA* apoplastic wash fluid spectra. However, a feature of [M+H]+ 472.2022 exhibiting the largest nominal reduction in intensity (160-fold) was undetectable in all Δ*lgsA* apoplastic wash fluid spectra (Figs. **3a**, **3c** and S**11a**). Targeted analysis revealed this metabolite could also be detected in whole-pseudostem extracts from WT-infected plants, though signal intensity was an order of magnitude lower than from apoplastic wash fluid samples. The functional genetic link between *lgsA* and this metabolite of 471.1949 Da (neutral mass) was confirmed by demonstrating that metabolite production could be restored by reintroduction of a WT *lgsA* sequence into both Δ*lgsA* strains (Fig. **3a**). The metabolite of 471.1949 Da was absent from apoplastic wash fluid samples from *L. perenne* infected with *E. festucae* var *lolii* strain CT.

**Fig. 3.**
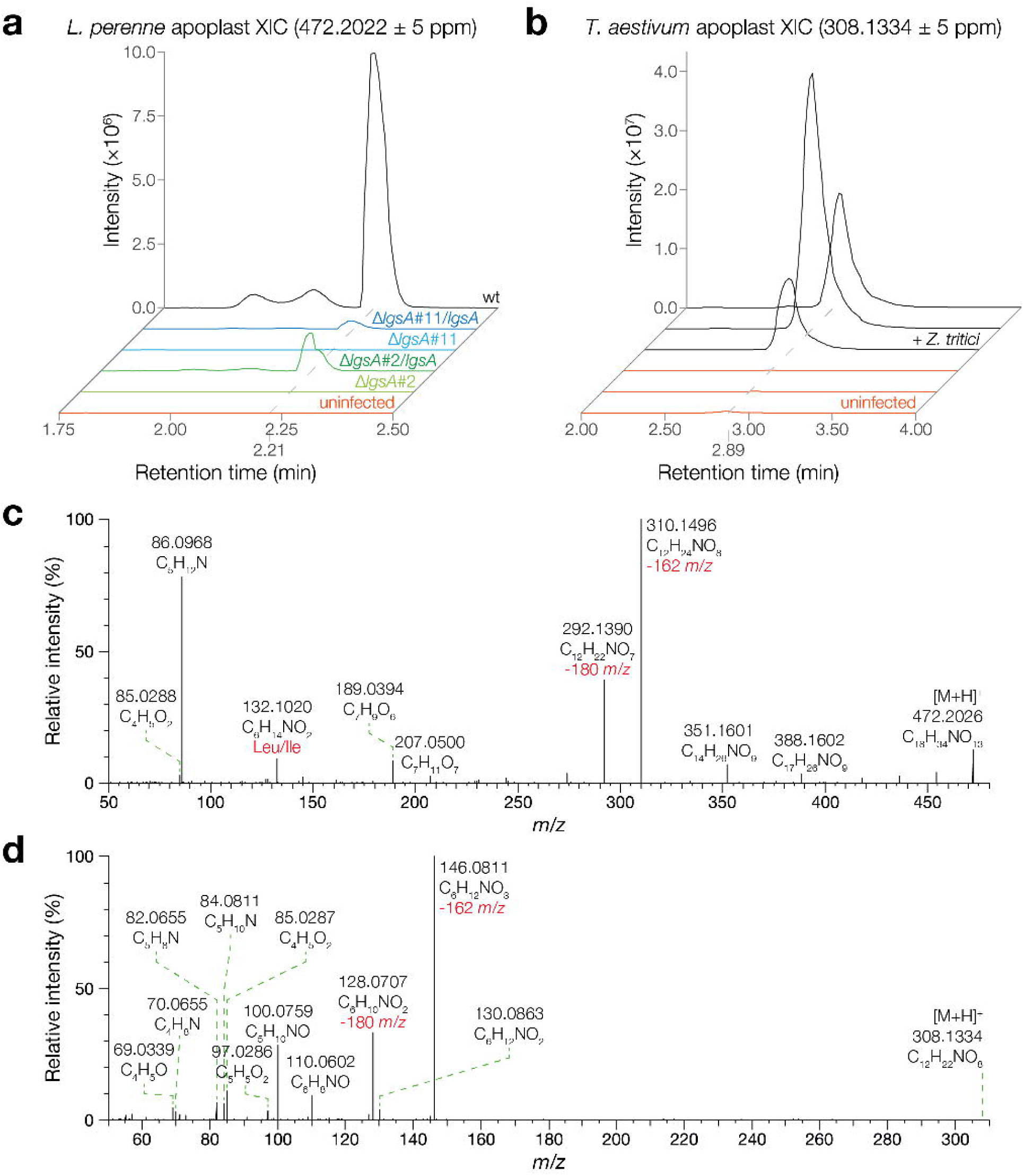
Identification of metabolites encoded by the *E. festucae* and *Z. tritici LGS* gene clusters. (a) Extracted ion chromatograms generated by LC-HR-MS analysis of apoplastic wash fluid samples extracted from uninfected, *E. festucae* Δ*lgsA*-infected (two independent mutants), *E. festucae* Δ*lgsA*/*lgsA*-infected (Δ*lgsA* strains complemented with *lgsA*), and *E. festucae* WT-infected *L. perenne* plants. Three plants were analysed per association, with representative examples shown here. (b) Extracted ion chromatograms generated by LC-HR-MS analysis of apoplastic wash fluid samples extracted from uninfected and *Z. tritici*-infected *T. aestivum* plants. Three plants were analysed per association, with results from all samples shown here. (c) High-resolution accurate mass positive ESI-HR-MSMS spectrum of the 472 *m*/*z E. festucae* LGS precursor ion fragmented by higher-energy collisional dissociation (HCD) at 20% energy. (d) High-resolution accurate mass positive ESI-HR-MSMS spectrum of the 308 *m*/*z* putative *Z. tritici* LGS precursor ion fragmented by HCD at 20% energy.

Integration and comparison of the ^13^C (*m/z* 473.2056) and ^18^O (*m/z* 474.2064) isotopologues abundance to the *m/z* 472.2022 base peak abundance established sum formula constraints of C_18-21_ and O_13-14_ for this LGS metabolite. The absence of a detectable ^15^N (*m*/*z* 473.1992) isotopologue and the even *m/z* of this compound suggested the presence of one nitrogen (Fig. **S12a**). The metabolite was therefore assigned the elemental composition of C_18_H_33_NO_13_ (calculated mass 471.1952 Da, observed mass 471.1949 Da). LC-HR-MSMS analysis suggested this LGS metabolite has a structure containing a leucine (Leu) or isoleucine (Ile) moiety shown by the diagnostic fragments of *m/z* 132.1020 (C_6_H_14_NO_2_) and *m/z* 86.0962 (C_5_H_12_N) (Fig. **3c**, Supporting Information Table **S8**). Neutral loss of one hexose moiety (162.053 Da (C_6_H_10_O_5_) or 180.0636 Da (C_6_H_12_O_6_)) from the precursor ion of *m/z* 472.2026 led to fragments of *m/z* 310.1496 (C_12_H_24_NO_8_) and *m/z* 292.1390 (C_12_H_22_NO_7_), respectively. A further neutral loss of 178.0476 Da (C_6_H_10_O_6_) is then observed, consistent with loss of an ester-linked hexuronic acid moiety, leaving the fragment of *m/z* 132.1020 (C_6_H_14_NO_2_), which represents the Leu/Ile-moiety (Fig. **3c**).

### Analysis of apoplastic wash fluid from *Zymoseptoria tritici*-infected wheat plants for LGS-like metabolites

To determine whether proteins similar to LgsA are encoded by genes in other fungi, a tBLASTn search was performed against the NCBI database using the *E. festucae* LgsA protein sequence as query. This analysis identified matches to NRPS-like proteins in *Zymoseptoria brevis* (60% amino acid identity), *Z. tritici* (60%), and *Ramularia collo-cygni* (68%) (Fig. **S7**) that share the same unusual T-C-A domain structure as LgsA. Homologues to other putative *LGS* cluster genes were either absent or located at different loci in these *Zymoseptoria* and *Ramularia* spp., with the exception of the putative NADP-binding oxidoreductase (*lgsB*), which was located immediately adjacent to the *lgsA* homologues of both *Zymoseptoria* species (Fig. **2**).

We therefore compared apoplastic wash fluid samples from wheat plants infected with *Z. tritici* to non-infected wheat plants by LC-HR-MS. These analyses showed that *Z. tritici*-infected wheat apoplastic wash fluid samples do not contain the 471 Da metabolite produced by the *E. festucae LGS* cluster. Data-dependent LC-HR-MSMS spectra generated from the *Z. tritici*-infected wheat apoplast samples were therefore filtered to identify metabolites exhibiting a neutral loss of 162.0523 Da (C_6_H_10_O_5_/hexose-moiety) or 178.0476 Da (C_6_H_10_O_6_/hexuronic acid-moiety, which are characteristic features of the *Epichoë* LGS product. Candidates exhibiting these neutral losses were eliminated if they were also present in the LC-HR-MS spectra of uninfected wheat apoplastic wash fluid samples, exhibited a substantially latter retention time compared to the 2.21 min *E. festucae* LGS metabolite, or exhibited elemental compositions or LC-HR-MSMS spectrum features that were not consistent with the properties expected of an LGS-like product. This left a single [M+H]^+^ 308.1334 feature with a retention time of 2.89 min as the sole remaining candidate for a *Z. tritici LGS* cluster product in this dataset, corresponding to a metabolite with an elemental composition of C_12_H_21_NO_8_ (calculated mass 307.1267 Da, observed mass 307.1261 Da) (Figs. **3b, 3d** and **S12b**). Analysis of the LC-HR-MSMS spectrum for this compound shows that the product ion of *m/z* 146.0811 remaining after the loss of the hexose-moiety as well as the corresponding fragments of *m/z* 128.0707 and 100.0759 could derive from a six-carbon amino acid, though this does not appear to be a Leu/Ile-moiety (Fig. **3d** and **S11b**). Given that the substrate-specifying “A-domain code” (Stachelhaus *et al*., 1999) of the *Z. tritici* LgsA homolog (DGLMYAVILK) has diverged somewhat from that of *E. festucae* LgsA (DALLYGIMAK) it is possible that these two proteins bind different amino acid substrates.

### LgsC is required for biosynthesis of the *E. festucae* LGS product

Given the *E. festucae LGS* cluster has a gene (*lgsC*; EfM3.062310) encoding a glycosyl transferase that is absent from the *Z. tritici* cluster, we hypothesised that this gene could encode an enzyme catalysing the conjugation of the second hexuronic acid present in the 471 Da *E. festucae* LGS product. We therefore prepared a replacement construct (pCE57) for *lgsC*, and transformed the linear product of this construct into protoplasts of *E. festucae*. A PCR screen of hygromycin-resistant transformants identified four transformants (Δ*lgsC*#6-6, #7-6, #11-4 and #12-6) in which the *lgsC* gene was deleted, with this result confirmed by Southern blot analysis (Fig. **S13**). Seedlings of *L. perenne* were infected with *E. festucae* WT, Δ*lgsC*#6-6 and Δ*lgsC*#12-6 and, as expected, *L. perenne* plants infected with the Δ*lgsC* mutant exhibited the same host interaction phenotype as WT-infected plants (Fig. **S14**). Whole-pseudostem extracts and apoplastic wash fluid samples were harvested from three mature plants of each association and analysed by LC-HR-MS. Apoplastic wash fluid samples from the Δ*lgsC* mutants lacked the 471 Da LGS product, demonstrating that LgsC is required for LGS biosynthesis in *E. festucae* (Fig. **4**). Targeted analysis revealed that Δ*lgsC*-infected material does not appear to accumulate any monoglycosylated LGS intermediates, suggesting that LgsC may act before LgsA in the LGS biosynthetic pathway.

**Fig. 4.**
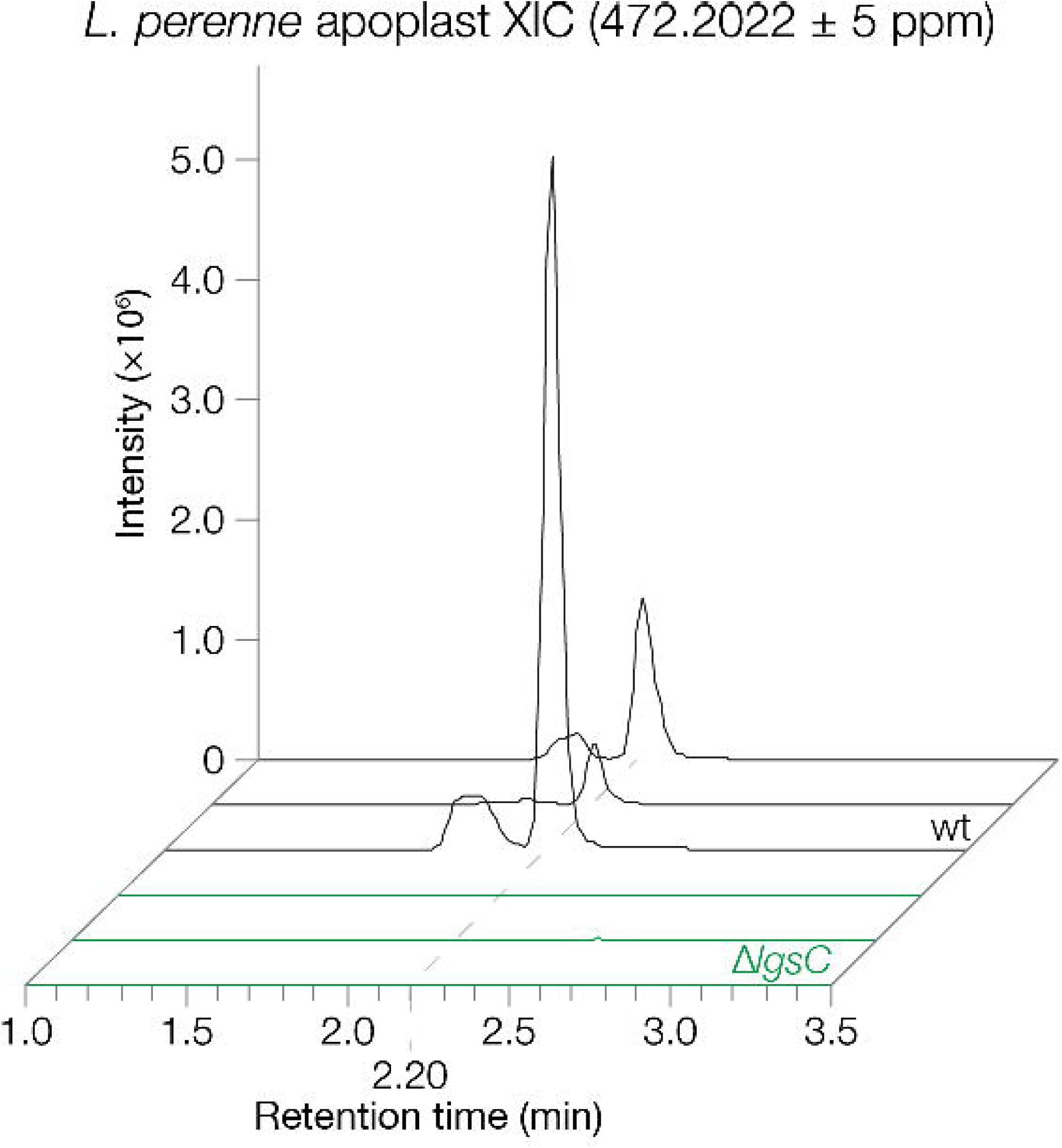
LgsC is required for synthesis of the *E. festucae* LGS product. Extracted ion chromatograms generated by LC-HR-MS analysis of apoplastic wash fluid samples extracted from *L. perenne* plants infected with *E. festucae* WT or Δ*lgsC* strains (two independent mutants). Three plants were analysed per association, with representative examples shown here.

## Discussion

Infection of *L. perenne* with the mutualistic fungal symbiont *E. festucae* results in a dramatic change to both the host and fungal transcriptome (Dupont *et al*., 2015; Winter *et al*., 2018), leading to a reprogramming of plant host and mycosymbiont metabolism. While there are many changes in primary metabolism (Rasmussen *et al*., 2008; Zhang *et al*., 2009; Dupont *et al*., 2015), it is the changes to fungal secondary metabolism that best define these associations as many compounds within this group have known bioprotective roles in natural ecosystems (Schardl *et al*., 2012). The preferential expression of genes encoding these alkaloid biosynthetic enzymes in the host plant makes it challenging to identify novel endophyte-induced metabolites given the complexity of the leaf extract metabolome. However, by focusing on the apoplast, where the endophyte grows, the metabolite complexity is reduced thereby enabling the identification of novel products. Here we describe for the first time a non-targeted analysis of the *L. perenne* apoplast metabolome and how it changes upon infection with *E. festucae*.

The metabolite differences between the apoplast metabolome of mock-infected versus *E. festucae* infected ryegrass is surprisingly uncomplex, with just 203 metabolite features identified distinguishing these two physiological states. This is in contrast to more complex metabolome changes reported for the plant fungal pathogen *V. longisporum* (Floerl *et al*., 2012). Of these features, 184 were shown to be enriched in the endophyte infected apoplastic fluid wash with the remaining 19 features corresponding to depletions in endophyte infected apoplastic fluid wash. As expected fungal metabolites known to be preferentially accumulated in the host are among those enriched upon endophyte infection, including peramine (Tanaka *et al*., 2005), and epichloëcyclins (Johnson *et al*., 2015). No lolines were detected in this analysis as the two *E. festucae* strains used lack this biosynthetic capability (Schardl *et al*., 2013).

The epichloëcyclins were among the most abundant metabolites synthesized in response to endophyte infection. These are RiPPs generated through proteolytic cleavage and cyclisation of linear peptide repeats that comprise the pro-peptide product of *gigA*. This gene is one of the most highly expressed in the endophyte-grass symbiotic interaction and is part of a four-gene cluster (Eaton *et al*., 2010). This cluster also includes *kexB*, which encodes a kexin presumably required for cleavage of the repeats from the pro-peptide; a gene encoding a protein with a DUF3328 domain, which is also found in UstYa/UstYb, which are proteins required for cyclisation of the ustiloxin precursors in *Aspergillus flavus* (Nagano *et al*., 2016; Ye *et al*., 2016); and a gene encoding a hypothetic protein of unknown function. The cyclic nonapeptides epichloëcyclin A, B and C, and the cyclic octapeptides epichloëcylin B and C (Johnson *et al*., 2015), were found in the apoplastic wash fluid of ryegrass infected with strain Fl1, whereas only epichloëcyclin A was found in apoplastic wash fluid from CT-infected plants. These results are in agreement with the repeat structures of the corresponding GigA peptides from these *E. festucae* strains. Both *E. festucae* strains have a functional copy of peramine synthetase, *perA*, and accumulate peramine in the apoplastic space as expected, given the known systemic distribution of this metabolite within the plant (Koulman *et al*., 2007). While metabolites with masses that match indole diterpenes known to be synthesised by these strains were detected (Saikia *et al*., 2012), the low signal intensity precluded their confirmation by LC-HR-MSMS. The absence of ergot alkaloids, which are known to be synthesized by these strains *in planta* (Chujo & Scott, 2014), is probably due to their lack of transport into the apoplast.

As anticipated, there were many metabolites of unknown structure, highlighting the utility of this type of analysis to identify novel endophyte-bioprotective metabolites. We therefore employed a combined genetics/metabolomics approach using the well-developed genetic system of *E. festucae* strain Fl1 (Scott *et al*., 2012; Scott *et al*., 2018) to demonstrate the utility of this apoplastic wash fluid method for identification of putative bioprotective fungal metabolites. For this analysis we chose to delete the NRPS-encoding gene *lgsA*, which is a component of a putative seven-gene SM cluster that is down regulated in *L. perenne* associations with three different symbiotically defective *E. festucae* mutants (Eaton *et al*., 2015). All seven genes were found here to be significantly upregulated in the Fl1 *in planta* transcriptome compared to axenic culture. As expected for a gene encoding a putative bioprotective metabolite (Tanaka *et al*., 2005; Young *et al*., 2005; Saikia *et al*., 2012), the axenic culture and plant interaction phenotypes of the *E. festucae* Δ*lgsA* mutant were indistinguishable from WT. A 471 Da metabolite that was absent in apoplastic wash fluid samples from plants infected with the Δ*lgsA* mutants was identified by LC-HR-MS in extracts of apoplastic wash fluid from *L. perenne* plants infected with WT *E. festucae* Fl1. The failure to identify this compound in an equivalent comparative analysis of whole-pseudostem extracts of these same plants highlights the power of working with the much less complex apoplastic metabolome. LC-HR-MSMS analysis suggests this LGS metabolite has a structure containing a Leu/Ile moiety linked to a glycoside constituent. Given the relatively small quantities of apoplastic wash fluid obtained using the method described here, definitive structural determination will likely require development of a high-volume extraction method or an alternative approach such as overexpressing the cluster genes in a heterologous fungal host to obtain sufficient product for NMR analysis (Van de Bittner *et al*., 2018; van Dolleweerd *et al*., 2018). The absence of this 471 Da metabolite in the apoplastic fluid of *L. perenne* with strain CT suggests that the genes are either absent, mutated, or not expressed in this strain. However, the presence of the *LGS* cluster across several *Epichloë* species suggests this is a widespread SM of this genus (Eaton *et al*., 2015).

A comparative synteny analysis of the genomes of other filamentous fungi identified highly conserved homologues to the NRPS-encoding *lgsA* gene in the genomes of the cereal pathogens *R. collo-cygni*, *Z. brevis*, and *Z. tritici*, with conserved homologues to the putative NADP-binding oxidoreductase-encoding gene *lgsB* located immediately adjacent in the *Zymoseptoria* genomes. LC-HR-MS analysis of extracts of apoplastic wash fluid samples from wheat infected with WT *Z. tritici* compared to apoplastic wash fluid from uninfected wheat plants identified a 307 Da metabolite that appears to have some structural similarities to the 471 Da metabolite, but with only a single glycosyl constituent. This structure is consistent with the absence of a conserved homologue to the putative glycosyl transferase-encoding gene *lgsC* in these species. However, more substantial evidence, such as analysis of a *Z. tritici* Δ*lgsA* mutant, will be required to prove the hypothesised link between the *Z. tritici LGS* cluster and this 307 Da metabolite.

Because *lgsA* and *lgsB* are conserved between the *LGS* clusters of both *Zymoseptoria* and *Epichloë* spp., we propose that a Leu/Ile substrate is selected and activated by the A-domain of LgsA, then tethered to the LgsA T-domain via a thioester bond. The LgsA C-domain would then catalyse ester bond formation between the thio-tethered Leu/Ile moiety and a hexose substrate that is selected and oxidised by LgsB into the corresponding hexuronic acid. LgsC would add the glycosyl group observed in *E. festucae*, either before or after the LgsA-catalysed condensation event. The final LGS pathway product would then be exported into the apoplast by a transporter, presumably the putative ABC transporter encoded by *lgsD* in *E. festucae*, and one of the putative MFS sugar transporter genes that are co-clustered with the *lgsA* homologue in *Zymoseptoria* species. While not specifically analysed here, other co-clustered genes in *Epichloë* and *Zymoseptoria* spp. may also encode proteins that contribute to LGS biosynthesis in their respective species.

While we do not know what the biological functions are for the compounds identified here, we make two observations: (i) the gene clusters that encode the enzymes for the synthesis of these novel SMs appear to be limited to filamentous fungi that infect monocots of the subfamily Pooideae, and (ii) this gene cluster is dispensable for a mutualistic symbiotic interaction between *E. festucae* and *L. perenne*. The absence of a host interaction phenotype for the *E. festucae* Δ*lgsA* mutant suggests the product has a bioprotection role. Alternatively, these metabolites may have a role in virulence through suppression of host immunity. It will be interesting to analyse the host interaction phenotype of a *Z. tritici* Δ*lgsA* mutant.

In conclusion, we describe for the first time how the metabolome of *L. perenne* changes upon infection with either asexual or sexual isolates of *E. festucae*. Interestingly, the changes are not as dramatic as observed for fungal pathogen infection by *V. longisporum*, though the total number of studies in this field are still rather limited. Given the lower complexity of the apoplast metabolome in comparison to the metabolome of green tissues we have demonstrated here the power of a combined genetics/metabolomics approach to identify new metabolites in host apoplastic wash fluids. It will be of considerable interest to determine the structures of other unknown metabolites in this and other fungal-plant associations, and to identify the biological function of these metabolites in these symbioses.

## Acknowledgments

This research was supported by a grant from the Tertiary Education Commission to the Bio-Protection Research Centre and by Massey University. BS was supported by an Alexander von Humboldt Research Award.. IF was supported by the Deutsche Forschungsgemeinschaft (ZUK 45/2010). The authors thank Matthew Savoian, Panimalar Vijayan and Niki Minards (Manawatu Microscopy and Imaging Centre) for technical assistance, Nazanin Noorifar (Massey Univesity) for assistance with plant inoculations, David Lun (Massey University) for assistance with MS, Richard Johnson (AgResearch) for provision of *E. festucae lgsA* sequences, Benedikt Ni (University of Goettingen) for compiling the Epichloë metabolite data base and Christopher Schardl (University of Kentucky) for provision of *Clavicipitaceae* genome sequences.

## Author contributions

KG, KF, DB, IF and BS planned and designed the research. KG, KF, DB, CE, AR, and PS performed the experiments. KG, KF, DB, CHM, IF and BS analysed the data. KG, DB and BS wrote the manuscript.

## Supporting Information

**Fig. S1** Principle component analysis (PCA) of 203 metabolite features.

**Fig. S2** Extracted ion chromatogram of peramine.

**Fig. S3** Extracted ion chromatogram of epichloëcyclins A-E.

**Fig. S4** Confirmation of the identity of epichloëcyclins A-E.

**Fig S5** *gigA* gene cluster organisation and sequences.

**Fig. S6** Extracted ion chromatogram of a putative peptide.

**Fig. S7** LgsA predicted domain structure and multiple sequence alignment.

**Fig. S8** l*g*sA deletion and complementation construct design and strain screening.

**Fig. S9** Culture morphology of WT and Δ*lgsA* strains.

**Fig. S10** Host interaction and cellular phenotypes of *L. perenne* infected with WT and Δ*lgsA* strains.

**Fig. S11** ESI-HR-MSMS fragmentation of 472 *m/z* metabolite in negative mode.

**Fig. S12** 472 and 308 metabolite isotope analysis.

**Fig. S13** Strategy for deletion of *E. festucae lgsC*, Southern and PCR analysis.

**Fig. S14** Host interaction phenotype of *L. perenne* infected with WT and Δ*lgsC* strains.

**Table S1** Biological material.

**Table S2** Primers used in this study.

**Table S3** Data matrix of raw data obtained by UPLC-ESI-TOF-MS-based metabolite fingerprinting analysis of the polar extraction phase, analysed in positive ESI-mode.

**Table S4** Data matrix of raw data obtained by UPLC-ESI-TOF-MS-based metabolite fingerprinting analysis of the polar extraction phase, analysed in negative ESI-mode.

**Table S5** Data matrix of raw data obtained by UPLC-ESI-TOF-MS-based metabolite fingerprinting analysis of the non-polar extraction phase, analysed in positive ESI-mode.

**Table S6** Data matrix of raw data obtained by UPLC-ESI-TOF-MS-based metabolite fingerprinting analysis of the non-polar extraction phase, analysed in negative ESI-mode.

**Table S7** Data matrix of 203 high quality metabolite features (false discovery rate < 0.003) obtained by metabolite fingerprinting (UPLC-ESI-TOF-MS analysis) of apoplastic wash fluids from mock-treated, FI1- and CT-infected *L. perenne*.

**Table S8** Infection markers identified by metabolite fingerprinting (UPLC-ESI-TOF-MS analysis) and verified by UHPLC-ESI-QTOF-HR-MSMS analysis or coelution.

**Table S9** *Epichloë* metabolite database.

**Table S10** Differences in expression of *lgs* cluster genes *in planta* compared to axenic culture.

