## Supplemental Data 1 for "Reprogramming of the apoplast metabolome of *Lolium perenne* upon infection with the mutualistic symbiont *Epichloë festucae*"

The following Supporting Information is available for this article:

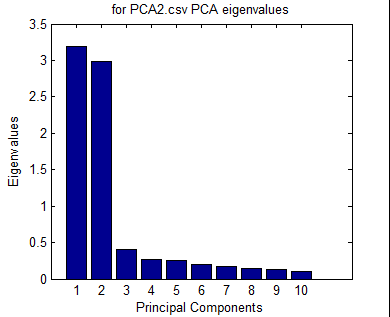

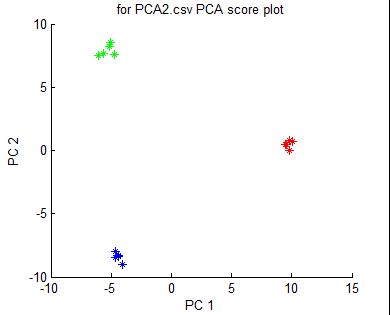

**a**

**b**

**Fig. S1** Principle component analysis (PCA) of 203 metabolite features with a false discovery rate of < 0.003, obtained by metabolite fingerprinting (UPLC-ESI-TOF-MS analysis) of apoplastic wash fluids from mock-treated (blue), or FI1 (red)- or CT (green)-infected *L. perenne*. (a) PCA scores plot and (b) PCA eigen values are shown. Five biological replicates per condition were used for the analysis.

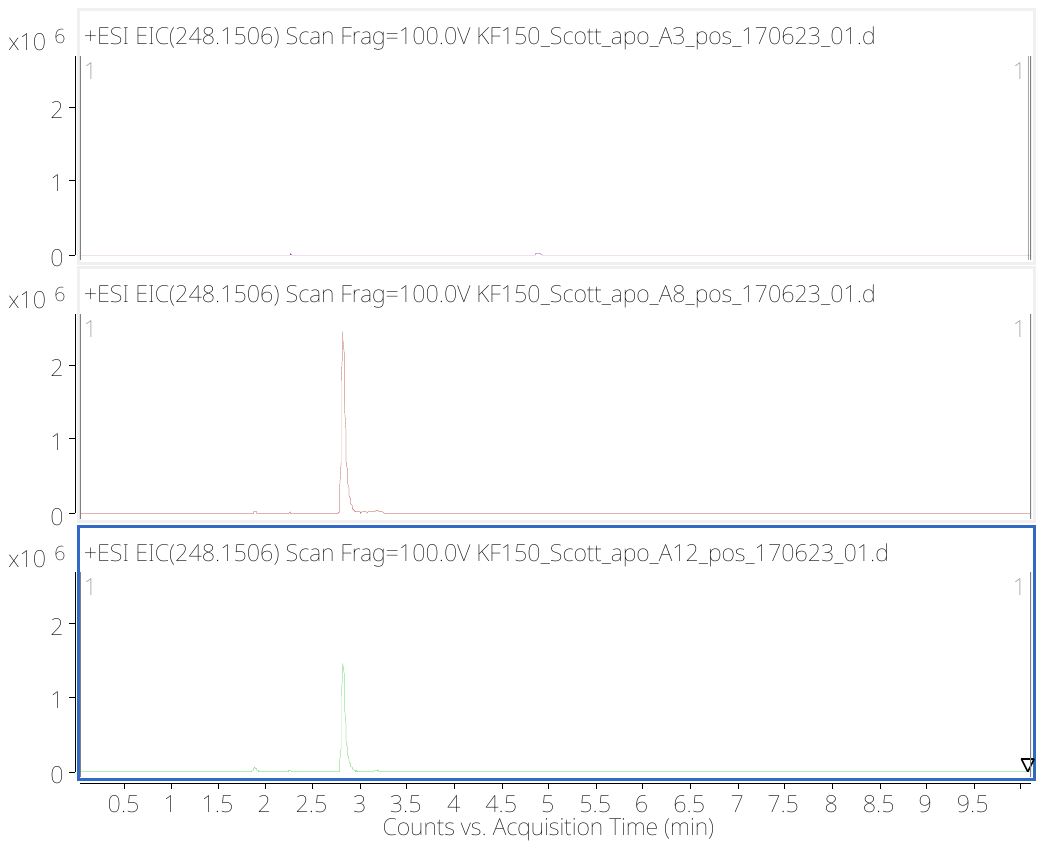

Mock

FI1

CT

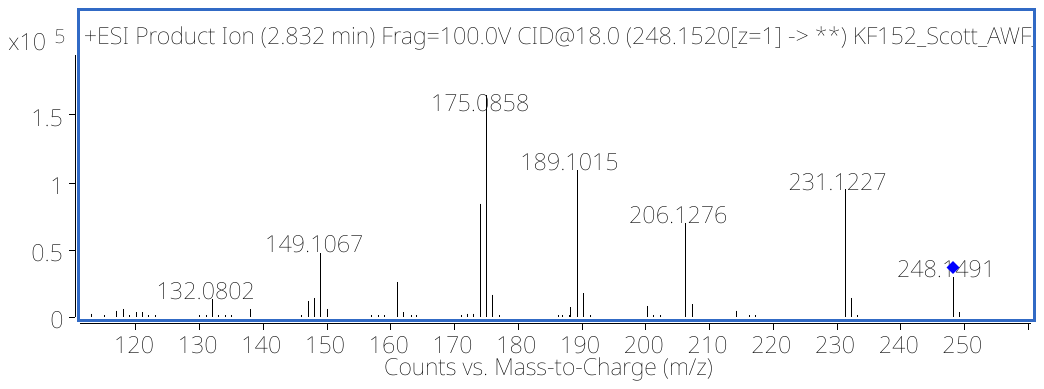

**a**

**b**

**Fig. S2** Extracted ion chromatogram of peramine ([M+H]^+^ 248.1506) in Fl1 and CT (a) and confirmation of the peramine structure by UHPLC-QTOF-MS/MS analysis. (b) Data are representative for five biological replicates per condition.

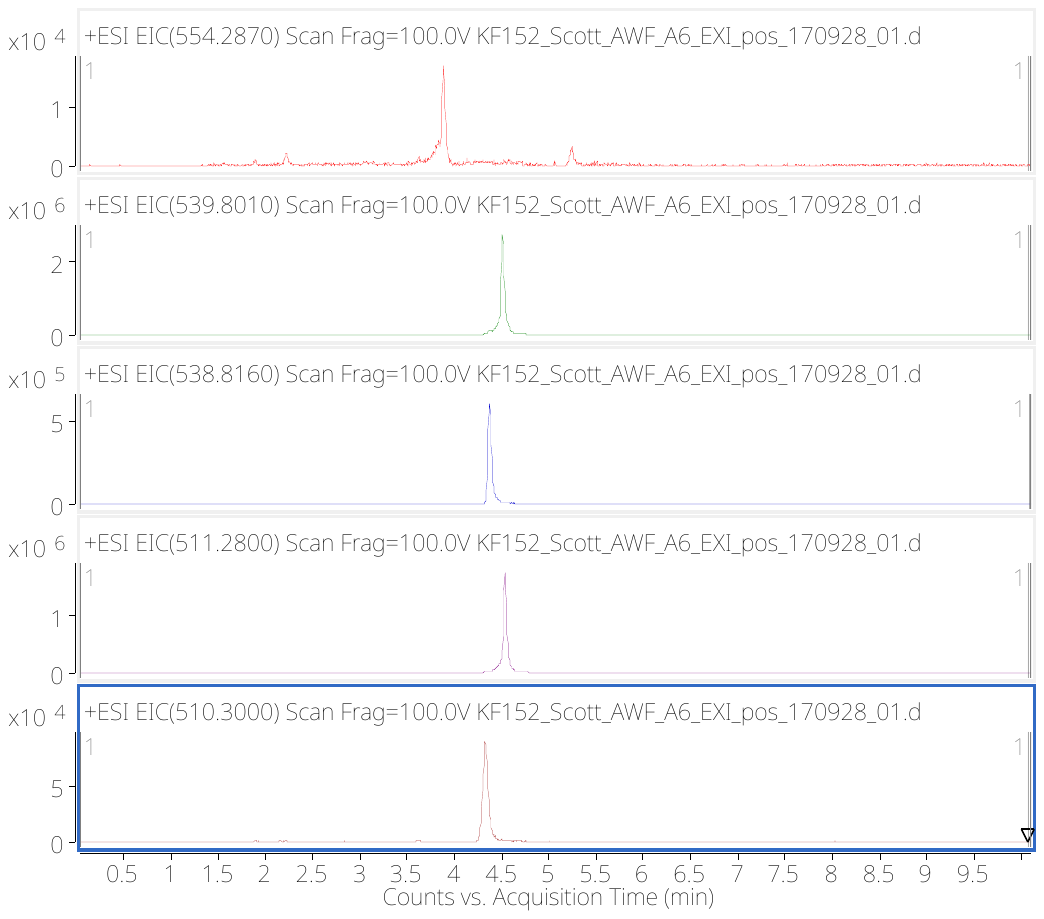

Epichloëcyclin A

Epichloëcyclin C

Epichloëcyclin D

Epichloëcyclin B

Epichloëcyclin E

**Fig. S3** Extracted ion chromatogram of epichloëcyclins A-E, obtained by UHPLC-QTOF-MS analysis of Fl1 samples. Data are representative for five biological replicates per condition.

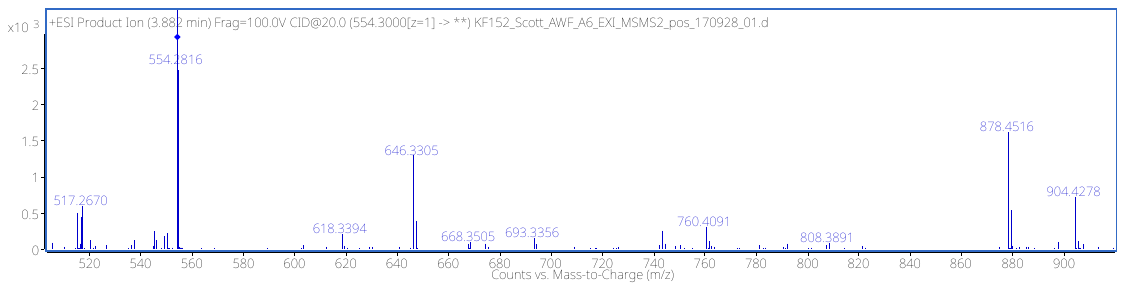

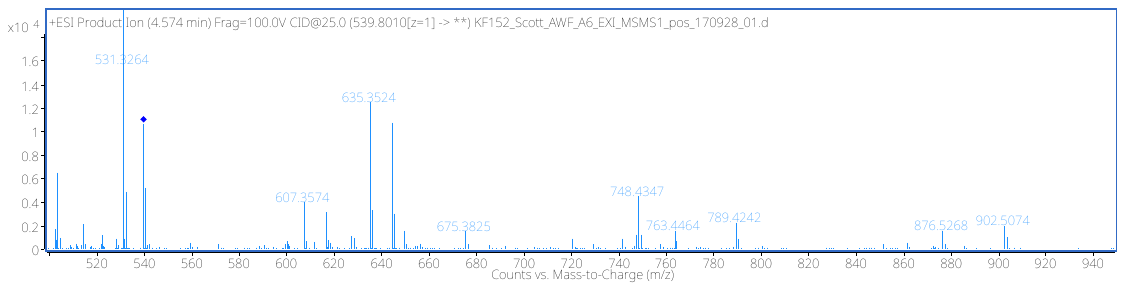

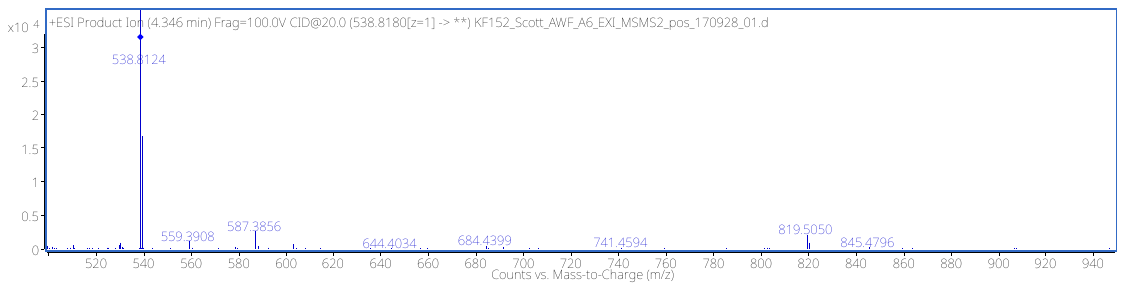

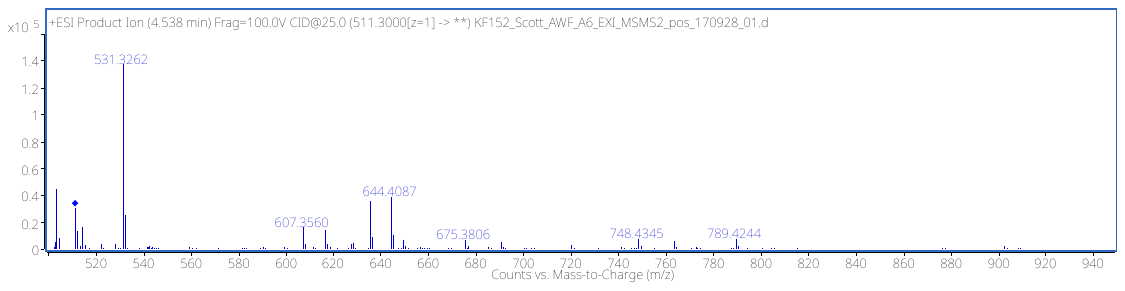

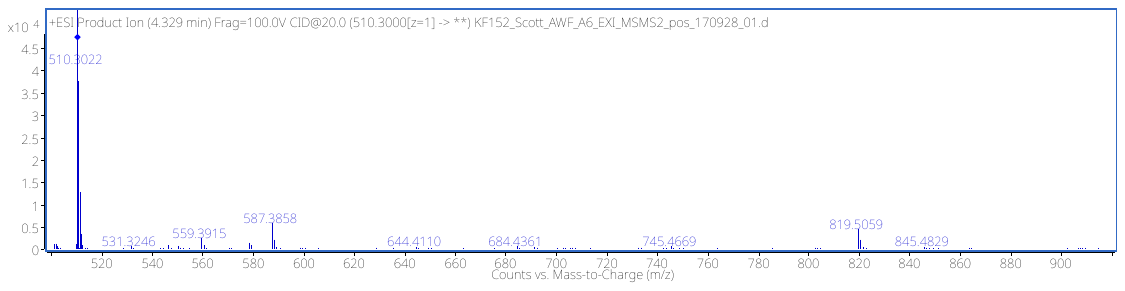

Epichloëcyclin A

Epichloëcyclin C

Epichloëcyclin D

Epichloëcyclin B

Epichloëcyclin E

**Fig. S4** Confirmation of the identity of epichloëcyclins A-E by UHPLC-QTOF-MS/MS analysis of the ions [2M+H]^2+^. The fragment spectra shown here for epichloëcyclins A-E match the spectra described by Johnson et al., 2015.

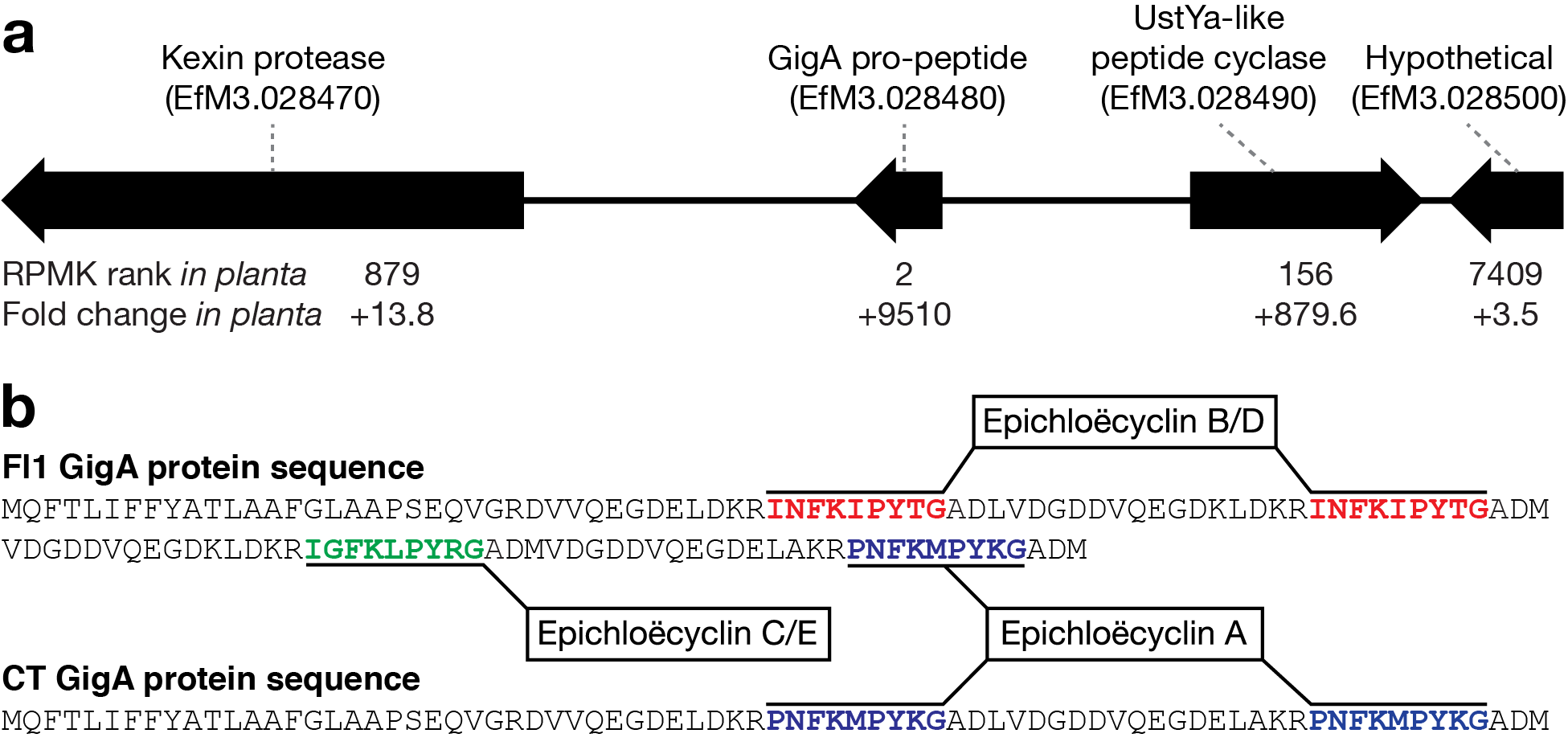

**Fig S5** *gigA* gene cluster and GigA protein sequences. (a) The *gigA* gene cluster annotated with gene expression level (ranked highest to lowest for all *E. festucae* genes expressed *in planta*) and fold change in expression *in planta* compared to axenic culture. The transcriptome data used here is available from the Sequence Read Archive (SRA) under Bioproject PRJNA447872. (b) GigA pro-peptide sequences from *E. festucae* strains Fl1 and CT annotated with the epichloëcyclin repeat sequences.

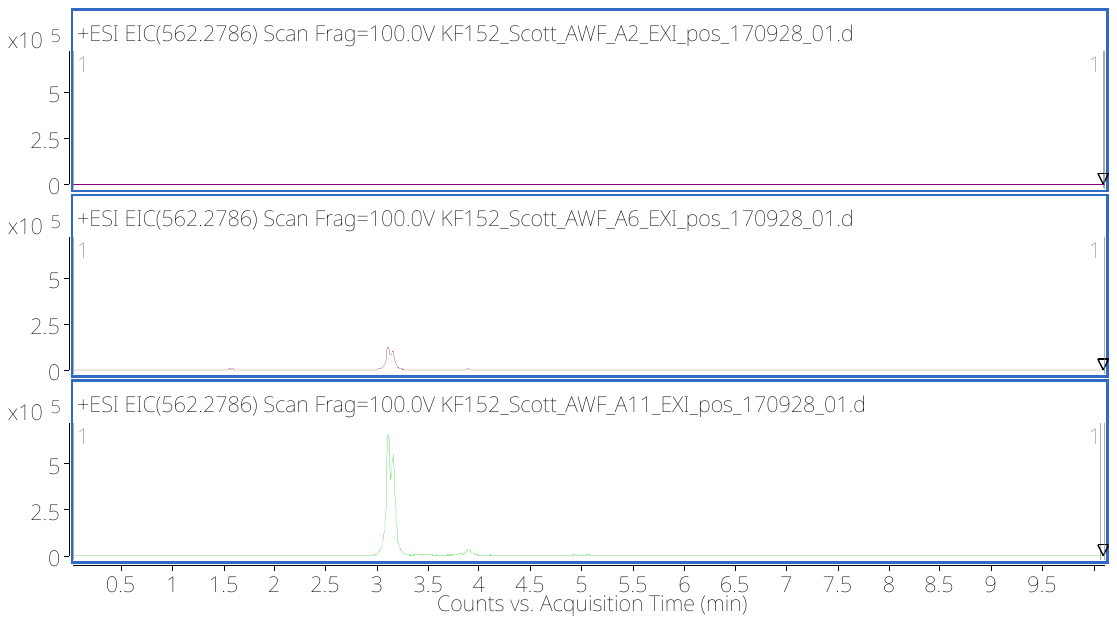

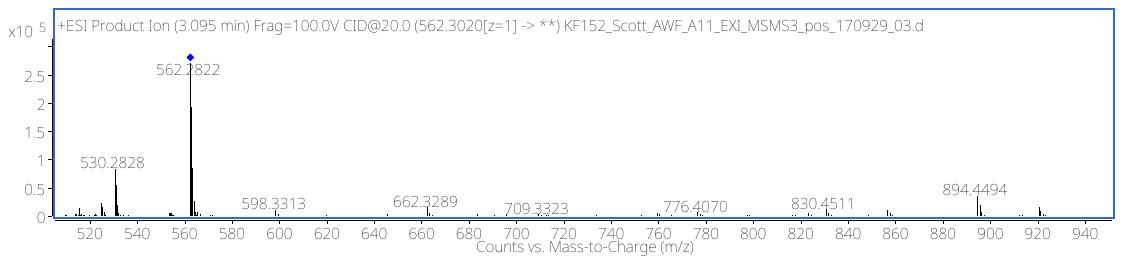

**a**

**b**

**Fig. S6**  Extracted ion chromatogram of a putative peptide of [2M+H]^2+^ 562.2786 (put. peptide 1), which is strongly enriched in CT. (a) Data are representative for five biological replicates per condition. (b) The fragment spectrum of this putative peptide 1 obtained by UHPLC-QTOF-MS/MS analysis is shown.

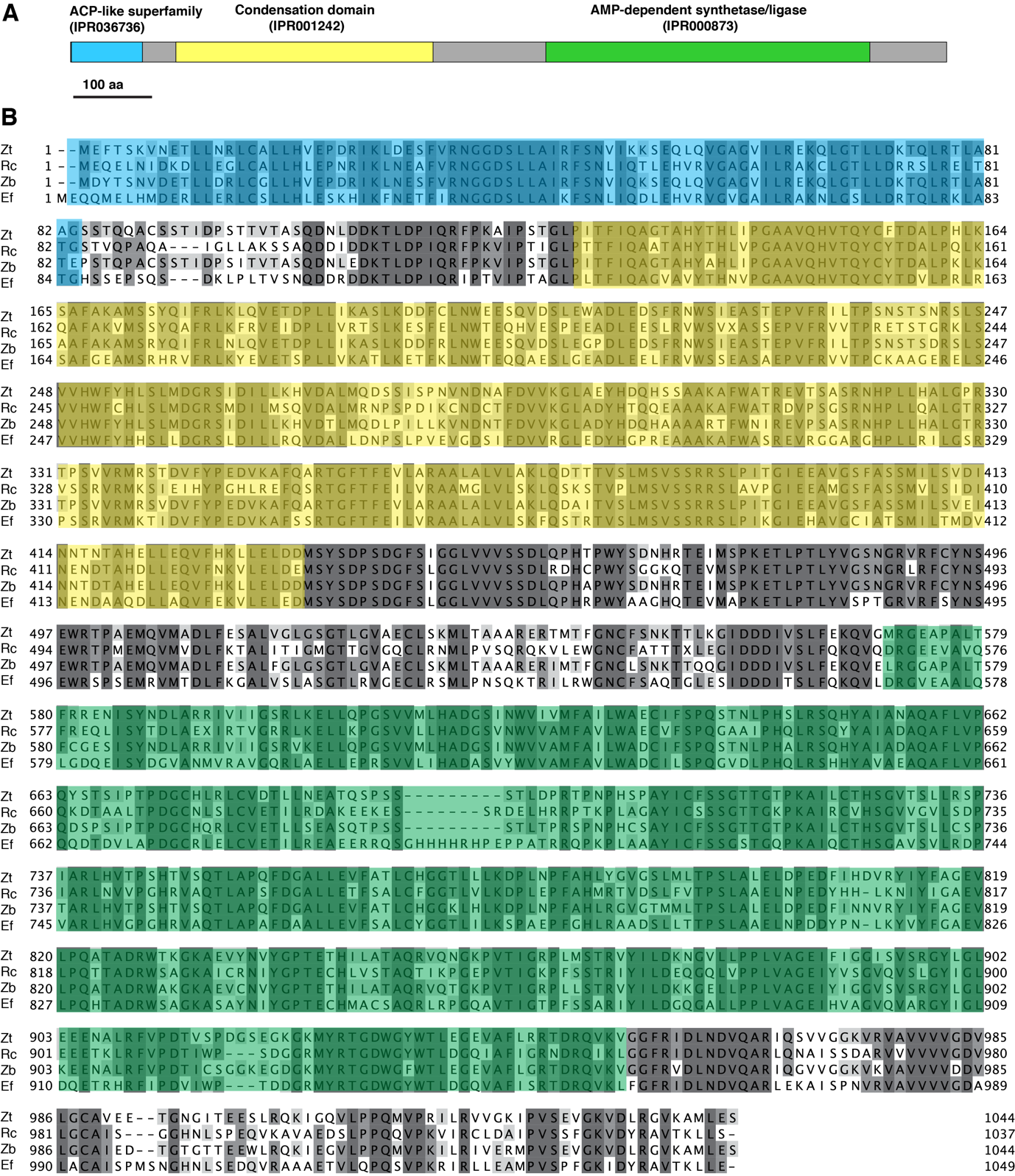

**Figure S7** LgsA predicted protein structure and multiple sequence alignment. (a) *Epichloë festucae* LgsA protein structure with respective InterproScan domains as indicated. Bar = 100 amino acids (aa). (b) Multiple sequence alignment of *E. festucae* (Ef) EfM3.056230, *Zymoseptoria tritici* (Zt) ZT1E4_G1886, *Ramularia collo-cygni* (Rc) RCC_03904, *Zymoseptoria brevis* (Zb) TI39_contig403g00012-encoded proteins. Domains as per (a), are indicated.

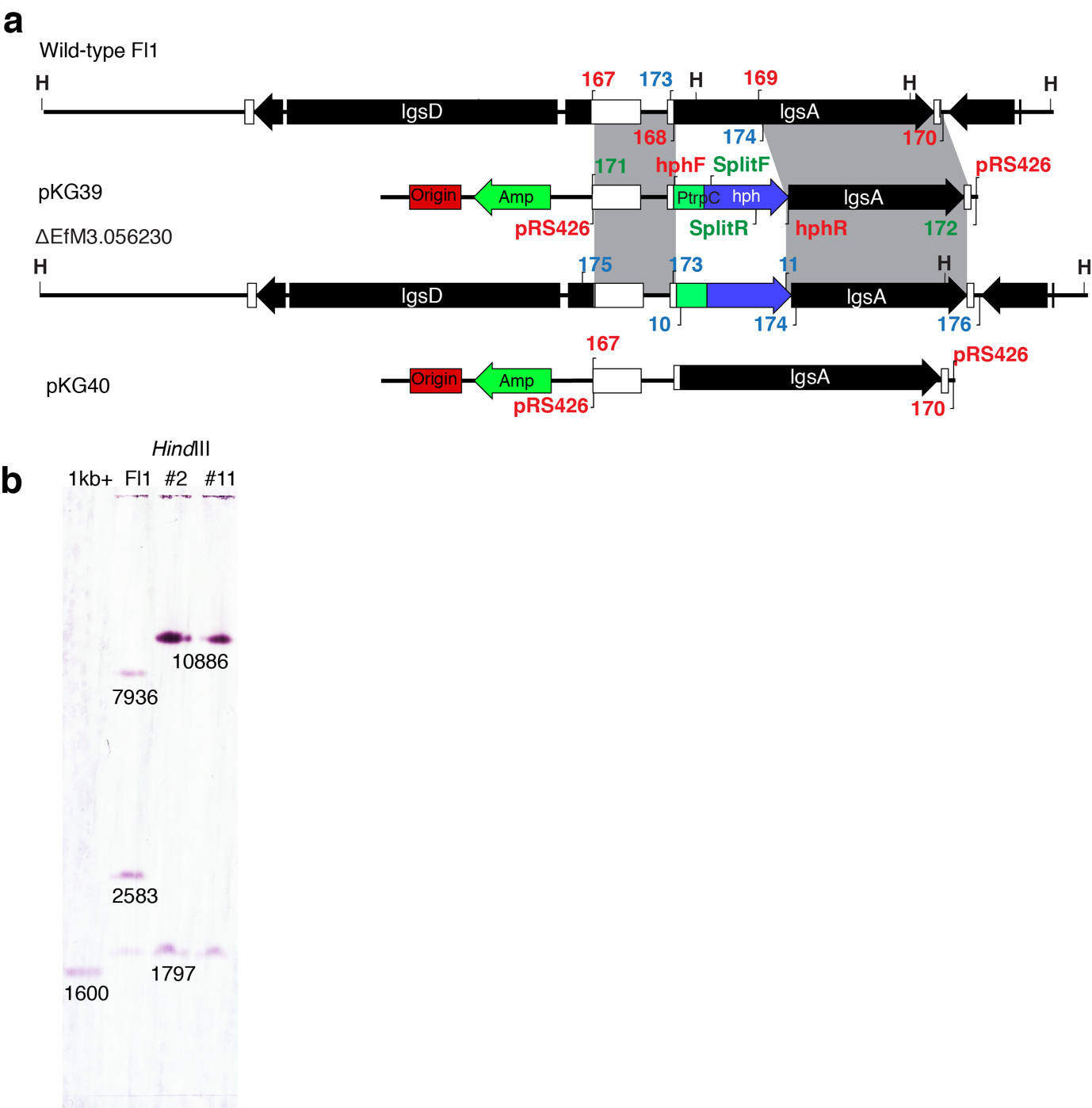

**Fig. S8** *lgsA* deletion and complementation construct design and strain screening. (a) Schematic of the WT (Fl1) *lgsA* genomic locus and linear insert of *lgsA* deletion construct pKG39 and complementation construct pKG40. Regions of recombination are indicated by grey shading. *Hin*dIII(H) restriction enzyme sites used for Southern analysis and PCR primers used for Gibson assembly (red), split marker transformations (green) and knock-out screening (blue) are as shown. (b) NBT/BCIP-stained Southern blot of *Hin*dIII genomic DNA digests (1.5 μg) probed with (DIG)-11-dUTP-labeled linear pKG39 purified PCR fragment (primers 171/172). Fragments of the expected size are as shown.

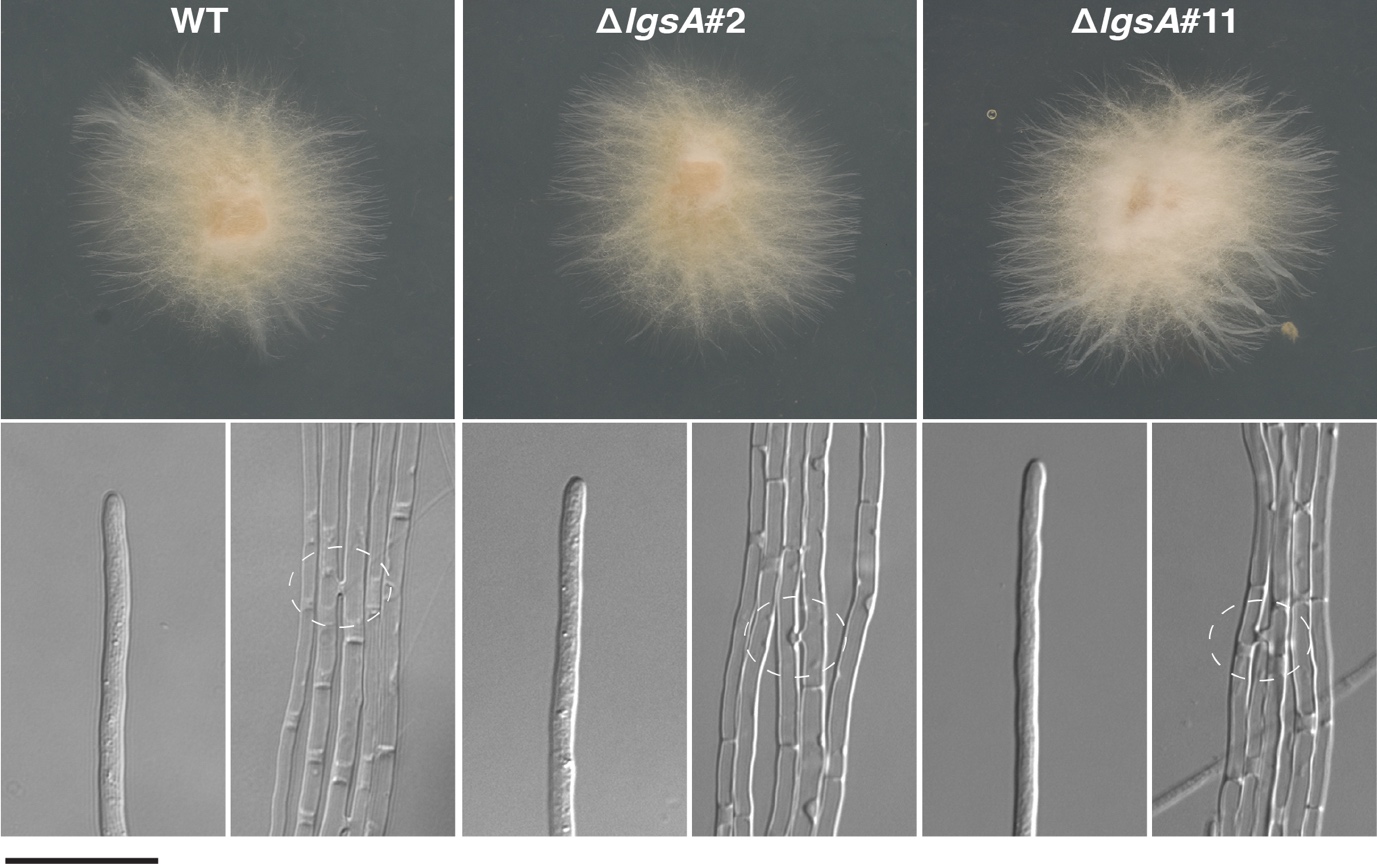

**Fig. S9** Culture morphology of WT and *∆lgsA* strains. (a) Culture morphology of WT and *∆lgsA* strains grown on PD agar for seven days at 22°C. (b) DIC images of hyphae undergoing cell-cell fusion on 1.5% water agar (highlighted by broken circles). Bar = 20 μm.

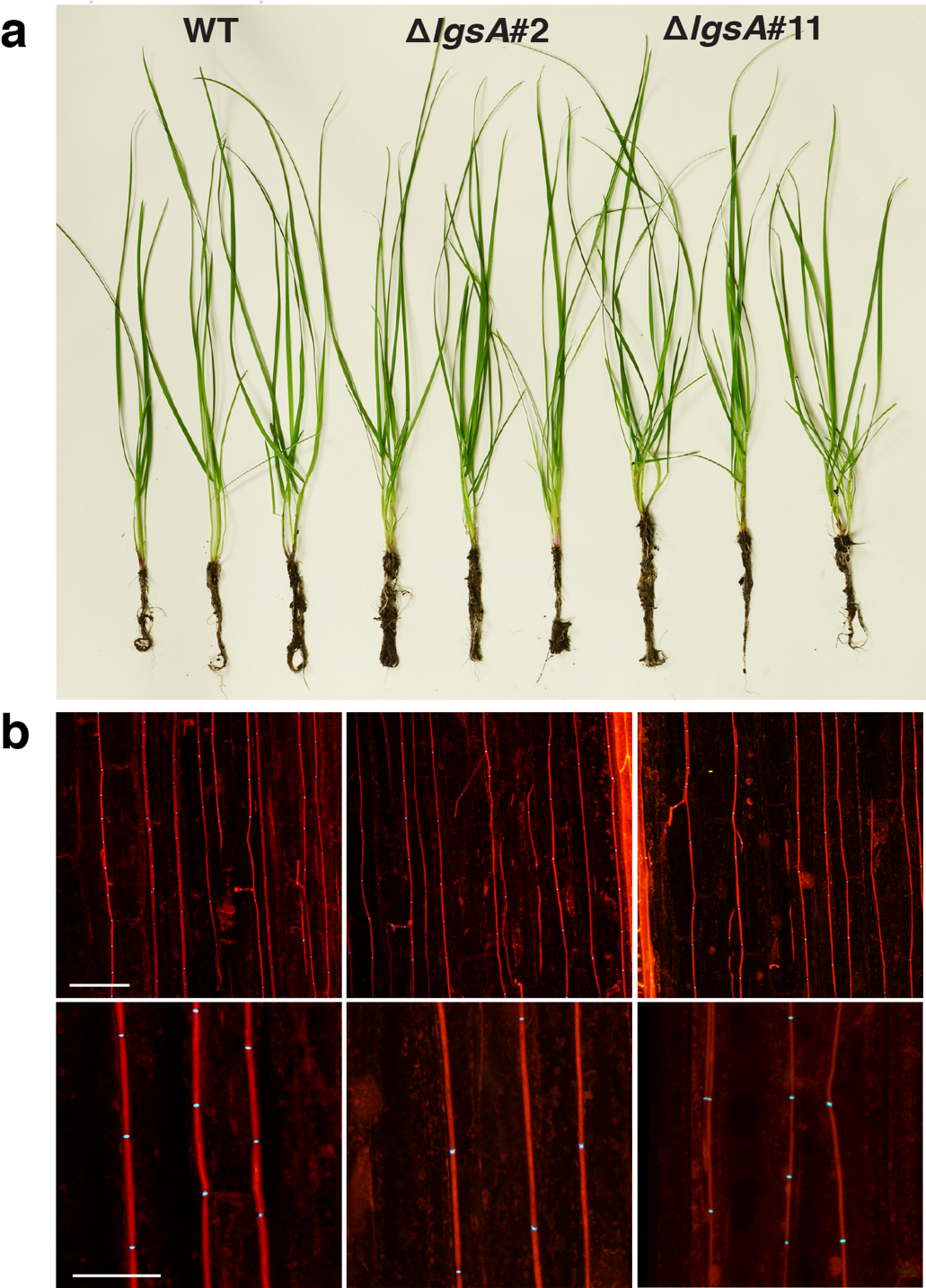

**Fig. S10** Host interaction and cellular phenotypes of *L. perenne* infected with WT and *∆lgsA* strains*.* (a) Host interaction phenotype of *L. perenne* plants infected with WT and *∆lgsA* strains at 8 weeks post planting. (b) Confocal depth series images of *L. perenne* leaf sheaths infected with WT and *∆lgsA* strains. Samples were stained with aniline blue (detects β-glucans; red) and WGA-AF488 (detects chitin; blue). Bar = 25 μm.

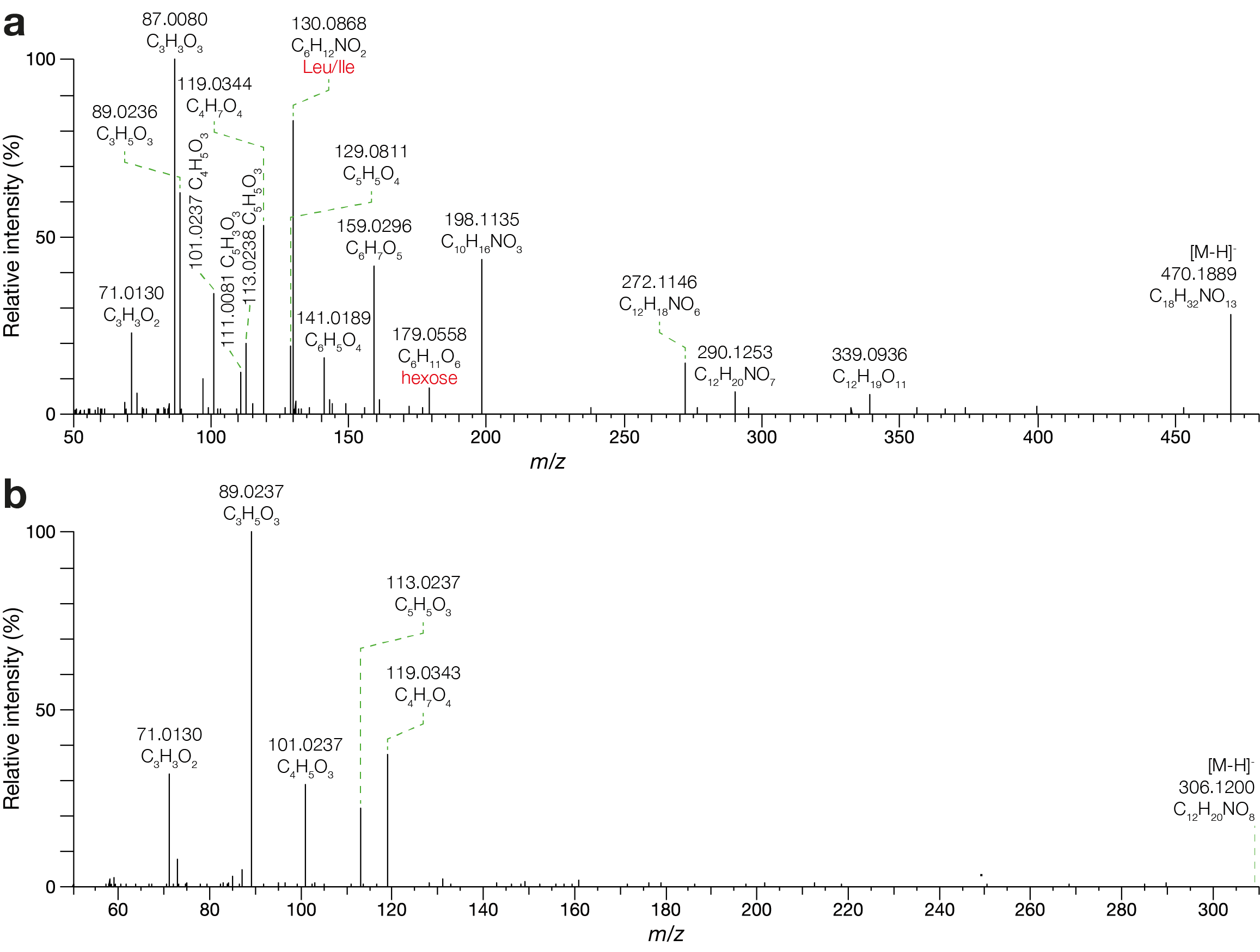

**Fig. S11** Negative mode ESI-MS/MS spectra. (a) High-resolution accurate mass negative ESI-MS/MS spectrum of the 470 *m*/*z* *E. festucae* LGS precursor ion fragmented by higher-energy collisional dissociation (HCD) at 20% energy. (b) High-resolution accurate mass negative ESI-MS/MS spectrum of the 306 *m*/*z* putative *Z. tritici* LGS precursor ion fragmented by HCD at 20% energy.

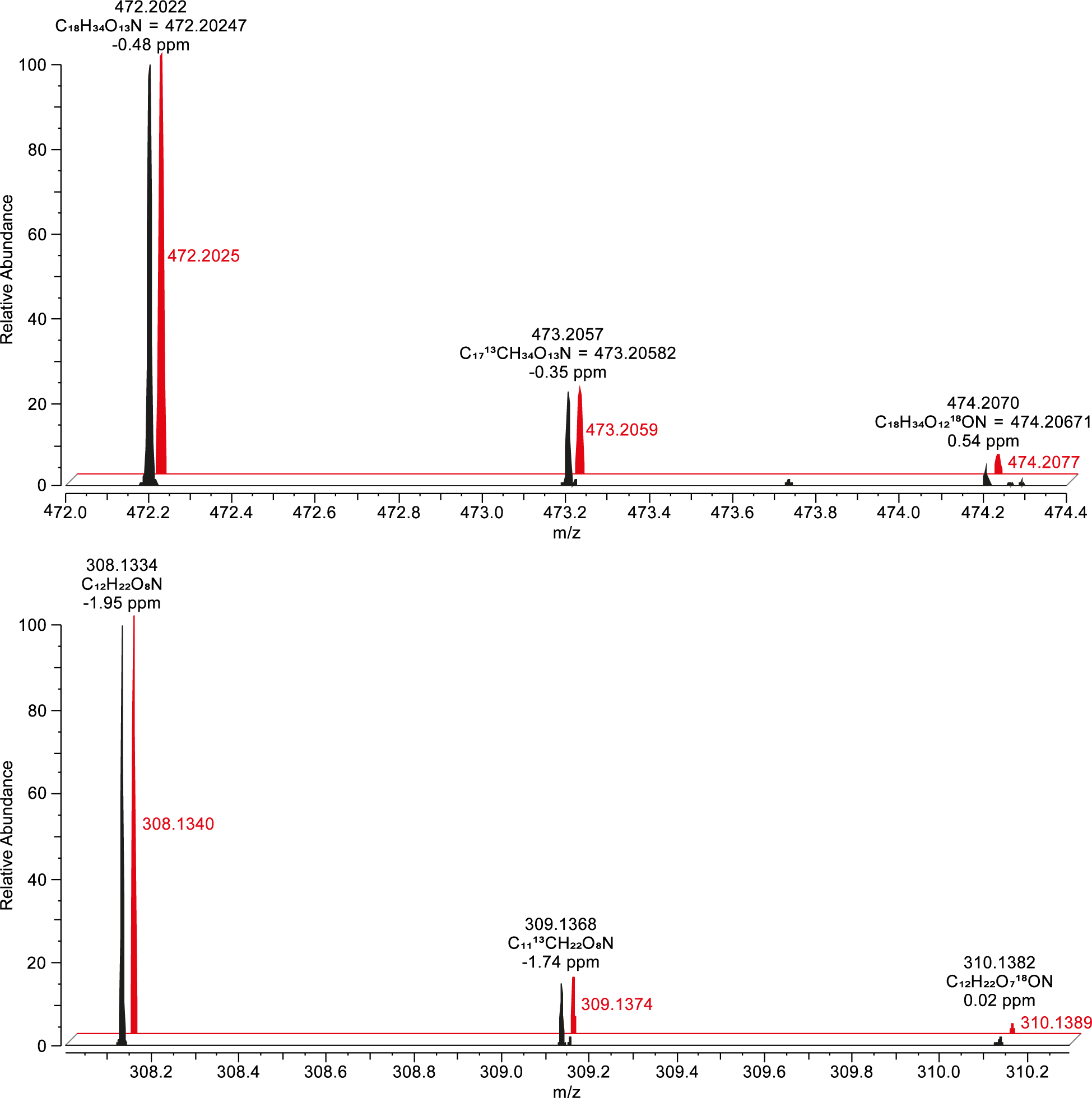

**Fig. S12** Observed (black) and simulated (red) isotope distributions for the 472 *m*/*z* (top) and 308 *m*/*z* (bottom) metabolites assuming molecular formulas of C_18_H_33_O_13_N + H^+^ and C_12_H_21_O_8_N + H^+^, respectively. Isotope distribution simulations were performed using Thermo FreeStyle 1.6.

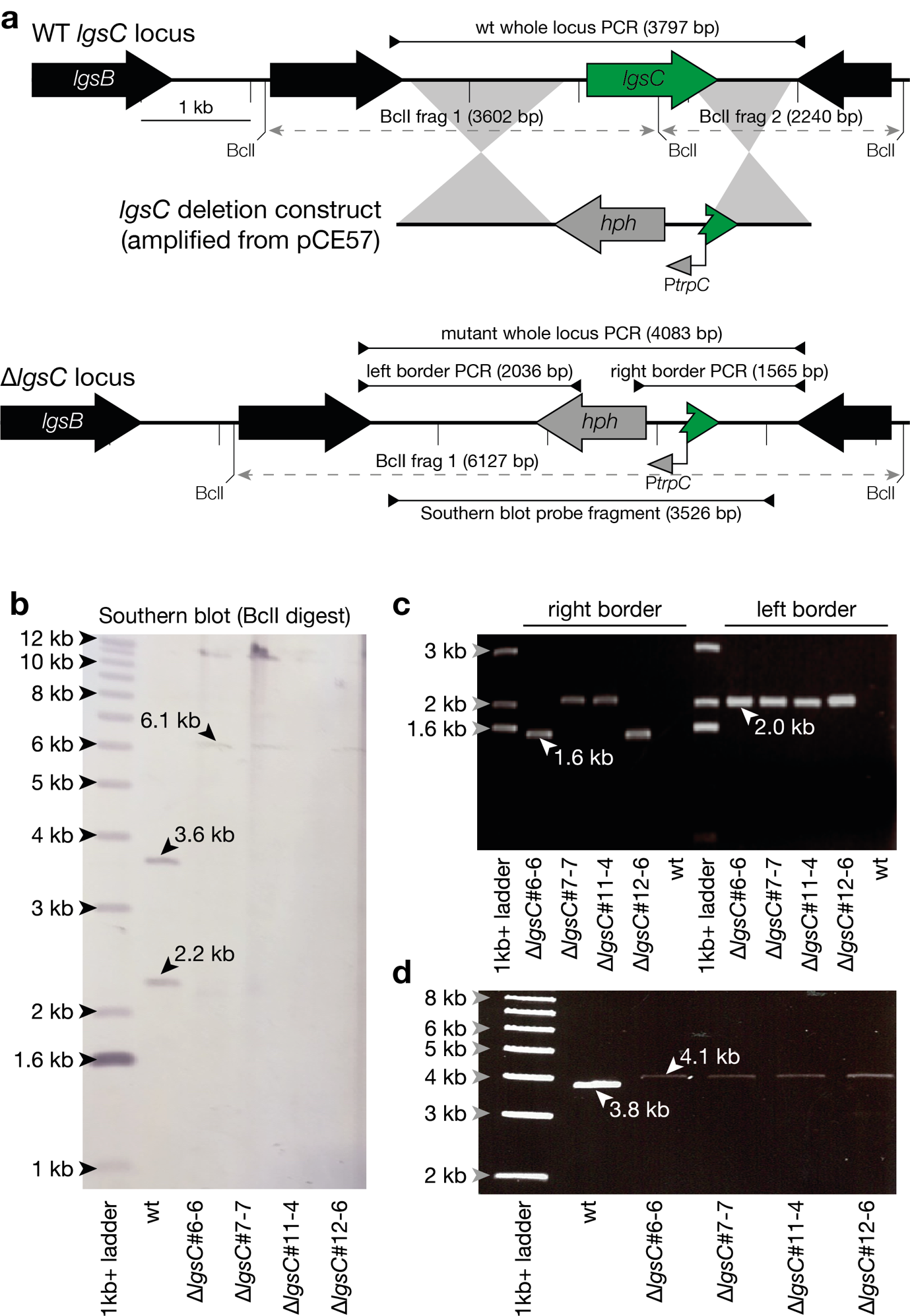

**Fig. S13** Strategy for deletion of *E. festucae lgsC*, Southern and PCR analysis. (a) Schematic of the *E. festucae* Fl1 (wild-type, WT) *lgsC* genomic locus, the linear *lgsC* deletion construct amplified from plasmid pCE57 using primers pRS426-cpsA-F and cpsA-pRS426-R, and the *∆lgsC* genomic locus. WT and mutant loci are annotated with *Bcl*I cut sites and amplified regions to aid interpretation of the Southern blot and PCR screening results, respectively. Homologous flanking regions enabling targeted recombination are indicated by grey shading. (b) NBT/BCIP-stained Southern blot of *Bcl*I-digested genomic DNA (1.5 μg per lane) probed with (DIG)-11-dUTP-labeled linear pCE57 purified PCR fragment (primers AR84/AR85). Fragments of the expected size are as shown. (c) Gel photo showing products of PCR amplification across the left (primers cps8/TC45) and right (TC44/cps7) flanking regions used to mediate integration at the target locus in four putative *∆lgsC* strains. No amplification of the WT control is expected. (d) Gel photo showing products of PCR amplification (primers cps7/cps8) across the entire *∆lgsC* integration site. Amplification of *lgsC* is expected for the WT control.

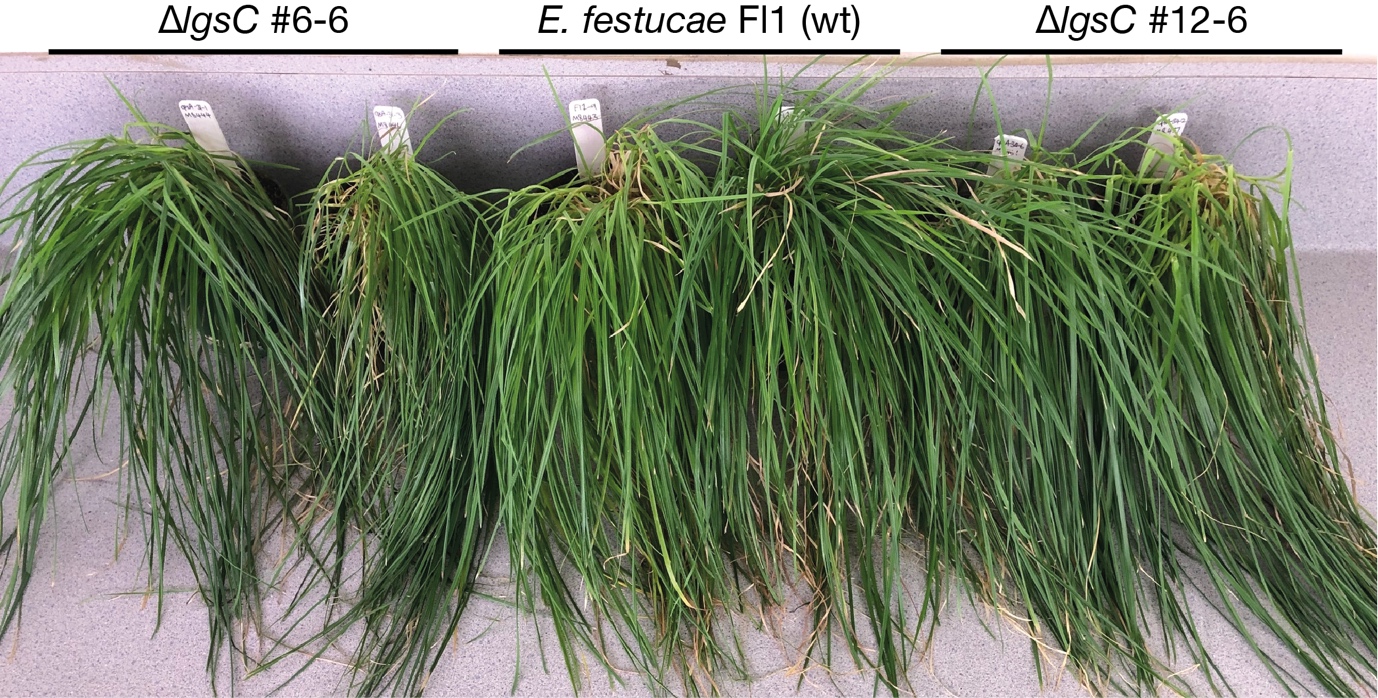

**Fig. S14** Host interaction phenotype of *L. perenne* infected with WT and *∆lgsC* strains*.* Host interaction phenotype of *L. perenne* plants infected with WT and *∆lgsC* strains #6-6 and #12-6 several months post-planting.

**Table S1** Biological material.

**Table S2** Primers used in this study.

**Table S3** Data matrix of raw data obtained by UPLC-ESI-TOF-MS-based metabolite fingerprinting analysis of the polar extraction phase, analysed in positive ESI-mode**.**

**Table S4** Data matrix of raw data obtained by UPLC-ESI-TOF-MS-based metabolite fingerprinting analysis of the polar extraction phase, analysed in negative ESI-mode.

**Table S5** Data matrix of raw data obtained by UPLC-ESI-TOF-MS-based metabolite fingerprinting analysis of the non-polar extraction phase, analysed in positive ESI-mode.

**Table S6** Data matrix of raw data obtained by UPLC-ESI-TOF-MS-based metabolite fingerprinting analysis of the non-polar extraction phase, analysed in negative ESI-mode.

**Table S7** Data matrix of 203 high quality metabolite features (false discovery rate < 0.003) obtained by metabolite fingerprinting (UPLC-ESI-TOF-MS analysis) of apoplastic wash fluids from mock-treated, FI1- or CT-infected *L. perenne*.

**Table S8** Infection markers identified by metabolite fingerprinting (UPLC-ESI-TOF-MS analysis) and verified by UHPLC-ESI-QTOF-MS/MS analysis or coelution.

**Table S9** *Epichloë* metabolite database

**Table S10** Differences in expression of *lgs* cluster genes *in planta* compared to axenic culture.
